# Jamaican fruit bats’ (*Artibeus jamaicensis*) competence for Ebola virus but not Marburg virus is driven by intrinsic differences in viral entry and IFN-I signaling antagonism

**DOI:** 10.1101/2024.10.17.618736

**Authors:** Sarah van Tol, Julia R. Port, Robert J. Fischer, Shane Gallogly, Trenton Bushmaker, Amanda Griffin, Jonathan E. Schulz, Aaron Carmody, Lara Myers, Daniel E. Crowley, Caylee A. Falvo, Jade C. Riopelle, Arthur Wickenhagen, Chad Clancy, Jamie Lovaglio, Carl Shaia, Greg Saturday, Jessica Prado-Smith, Yi He, Justin Lack, Craig Martens, Sarah L. Anzick, Lon V. Kendall, Tony Schountz, Raina K. Plowright, Andrea Marzi, Vincent J. Munster

## Abstract

Ebola virus (EBOV) and Marburg virus (MARV) are zoonotic filoviruses that cause hemorrhagic fever in humans. Bat species in both Chiropteran suborders host filoviruses, suggesting that bats may have coevolved with this viral family. Correlative data implicate bats as natural EBOV hosts, but neither a full-length genome nor an EBOV isolate has been found in any bats sampled. Here, we modelled filovirus infection in the Jamaican fruit bat (JFB), *Artibeus jamaicensis*. Bats were inoculated with either EBOV or MARV through a combination of oral, intranasal, and subcutaneous routes. EBOV-infected bats supported systemic virus replication and shed infectious virus orally. In contrast, MARV replicated only transiently and was not shed. *In vitro*, JFB cells replicate EBOV more efficiently than MARV, and MARV infection induced innate antiviral responses that EBOV efficiently suppressed. Experiments using VSV pseudoparticles or replicating VSV expressing the EBOV or MARV glycoprotein demonstrated an advantage for EBOV entry and replication early, respectively, in JFB cells. Overall, this study describes filovirus species-specific phenotypes for both JFB and their cells.

## Introduction

Bats are the presumed reservoir hosts for viruses of five filovirus genera. Extensive experimental infection [1–5] and epidemiological evidence [6–8] supports that the Egyptian rousette bat (ERB) (*Rousettus aegyptiacus*) is the natural host of the orthomarburgviruses, Marburg virus (MARV) and Ravn virus (RAVV). ERBs cave roost in large groups and antibodies against orthomarburgviruses are consistently found in several ERB populations [6–11]. Infectious virus has been found mainly in the saliva [1–3], but virus is also detected in urine and feces at lower levels [3]. Biting, social grooming, contaminated food (saliva), or aerosolized urine or feces could facilitate transmission although the natural route(s) remain unconfirmed [12]. Dehong virus, the newest member of the filovirus genus *Delovirus*, was isolated from Leschenault’s rousette (*R. leschenaultii*) lung samples in China [13]. A serological survey and tissue sampling of bats in Yunnan, China suggests Dehong virus circulates within the bat population. Lloviu virus (LLOV), a *Cuevavirus*, was isolated from European bats [14–18]. Initial detection of LLOV in Schreiber’s bats (*Miniopterus schrebersii*) was linked with die-offs [15], but no causal link between LLOV infection and death was established.

In addition to virus isolation, full genomes of other filoviruses have been detected in bats. The full-length genome of the *Dianlovirus* Mĕnglá virus (MLAV) was sequenced from *Rousettus spp.* in China [19]. In 2018, the orthoebolavirus Bombali virus (BOMV) was identified in Angolan (*Mops condylurus*) and little free-tailed (*Chaerephon pumilus*) bats in Sierra Leone [20]. Since BOMV’s identification, molecular and serological data indicate an extensive distribution within Africa [21–25].

Aside from BOMV, the molecular evidence for orthoebolaviruses naturally circulating in bats is sparse [26]. Serological data support that several bat species, including the hammer-headed fruit bat (*Hypsignathus monstrosus*), are exposed to filoviruses [9, 27]. Oppositionally, a study that evaluated viral molecular signatures of reservoir hosts predicted that Bundibugyo virus (BDBV) and Taï Forest virus (TAFV) circulate in a primate host [28]. The lack of molecular evidence for orthoeboloviruses in bats, aside from BOMV, suggests that either the incorrect bat species are being sampled or a non-bat host may be the natural host unlike the other bat-associated filoviruses [29].

Chiroptera, the mammalian order of bats, includes over 1,400 species. Bats are diverse and occupy various niches and specialize in different diets and behaviors. The two bat suborders, Yinpterochiroptera and Yangochiroptera, diverged approximately 63 million years ago [30]. Filoviruses successfully infect bats from both suborders which suggests co-evolution between bats and filoviruses. The factors that predict which bat species could support replication and transmission of the different bat-associated filoviruses are unknown. For example, ERBs support disseminated replication and shedding of MARV, whereas infection with the various orthoebolaviruses is limited and does not uniformly induce an antibody response [2]. Additionally, the physiological and environmental stressors that regulate filovirus shedding intensity and transmission remain largely unknown.

Due to gaps in knowledge regarding within host parameters that dictate filovirus-bat species compatibility, we evaluated the potential of Jamaican fruit bats (JFB) (*Artibeus jamaicensis*) to support Ebola virus (EBOV) and MARV replication. We found that JFBs supported non-lethal, disseminated EBOV infection with minimal-to-mild clinical signs and histopathologic changes and shed infectious virus. In contrast, MARV replication was limited primarily to the inoculation site, no MARV RNA is detected in swabs, and an antibody response was not elicited uniformly. The difference between EBOV and MARV replication in live bats correlated with in vitro data that showed EBOV was more efficient than MARV in JFB cell entry and antagonism of the type I interferon (IFN-I) response. These differences between EBOV and MARV may aid in the identification of key filovirus-host interactions that are required for complementarity. Further, JFB can be used as an EBOV infection model to study the physiological and environmental stressors that impact infection kinetics and viral shedding.

## Methods

### Ethics statement

All animal experiments were conducted in an AAALAC International-accredited facility, approved by the Rocky Mountain Laboratories (RML) Animal Care and Use Committee and adhered to the guidelines put forth in the Guide for the Care and Use of Laboratory Animals 8th edition, the Animal Welfare Act, United States Department of Agriculture and the United States Public Health Service Policy on the Humane Care and Use of Laboratory Animals. Work with infectious EBOV and MARV under Biosafety level 4 (BSL4) conditions was approved by the Institutional Biosafety Committee (IBC) and conducted in RML’s BSL4 facility. For the removal of specimen from BSL4, virus inactivation of all samples was performed according to IBC-approved standard operating procedures (SOPs).

### Viruses

EBOV-Mayinga (NC_002549) and MARV-Ozolin (AY358025.2) were used for infection studies and *in vitro* experiments. EBOV-Kikwit (KT582109.1) and EBOV-Makona C05 (KP096420.1) as well as MARV-Angola (KY047763.1) and MARV-Musoke (DQ217792.2) were also used for *in vitro* studies. Replication-competent recombinant vesicular stomatitis viruses encoding a green fluorescent protein reporter gene (VSV-GFP) were used for *in vitro* studies. VSVwt-GFP and VSV-EBOVmayinga-GFP have been described previously [31]. The VSV-MARVozolin-GFP was generated following two cloning steps. First, the MARV-Ozolin glycoprotein (GP) gene (AY358025.2) was cloned into the VSV backbone as previously described for other filovirus GPs [31]. Next, the GFP gene was added as an additional ORF between the MARV-Ozolin GP and the VSV-L genes. The virus was recovered from plasmid following established protocols previously described [32]. Protein expression in the supernatant was verified by western blot analysis using antibodies against MARV GP (clone 127-8, 1:1,000; [33]) and VSV-M (23H12, 1:1,000; Kerafast, Inc.).

Virus was propagated in Vero E6 cells (ATCC CRL-1586) in DMEM supplemented with 2% fetal bovine serum (FBS), 1 mM L-glutamine, 50 U/mL penicillin, and 50 μg/mL streptomycin (DMEM2). Vero E6 cells were maintained in DMEM supplemented with 10% FBS, 1 mM L-glutamine, 50 U/mL penicillin, and 50 μg/mL streptomycin (DMEM10). No *Mycoplasma* nor contaminants were detected in any of the virus stocks.

### Cells

JFB, or *Artibeus jamaicensis* (Aj), cells including immortalized and primary kidney(Aji and AjKi_RML2 respectively), lung (AjLu_RML3), and uropatagium derived fibroblasts (AjUFi_RML6) and ERB, or *Rousettues aegyptiacus* (Ra), cells including primary kidney (RASKM), and primary lung (RaLu1.4) [34] were maintained in DMEM/F-12 supplemented with 1X MEM non-essential amino acids (NEAA) (Gibco), 10% FBS, 50 U/mL penicillin, and 50 μg/mL streptomycin (10DMEM/F-12). Vero E6, immortalized ERB kidney (RoNi), and immortalized human hepatoma (Huh-7) cells were maintained in DMEM10. All cells were maintained at 37°C, 5% CO_2_ and were *Mycoplasma* negative.

### Bat inoculation

Jamaican fruit bats from a closed colony housed at Colorado State University were used in this study. Upon arrival at RML, Bats were co-housed in same sex groups of up to six animals per cage. The study comprised 28 animals total, 14 males and 14 females. Bats were allowed to acclimate to the facility for five days before collecting samples. Bats were intranasally, orally, and subcutaneously inoculated with a total of 3.0 x 10^4^ TCID_50_ of EBOV-Mayinga (six females (EBOV01-06) and six males (EBOV07-12)) or MARV-Ozolin (six females (MARV01-06) and six males (MARV07-12)). Four bats (two males, two females) that arrived with the same shipment of bats were used as healthy controls for histologic, B cell receptor (BCR) sequencing, and gene expression analyses. All inoculations and subsequent manipulations were performed under isoflurane anesthesia. Four EBOV- and four MARV-inoculated bats were euthanized at 3-, 7-, and 28-days post-inoculation (DPI) to assess viral replication and host gene expression in tissue samples. Oropharyngeal and rectal swabs were collected to monitor viral shedding prior to infection, every two days after infection through 14-DPI, 21-DPI, and at necropsies. Blood was collected at baseline, on necropsy days, and on 14 and 21 DPI. At necropsy, whole blood was used for hematology and serum was used for clinical chemistry. Hematology analysis was completed on a ProCyte DX (IDEXX Laboratories, Westbrook, Maine). Serum chemistries were completed on a VetScan VS2 Chemistry Analyzer (Abaxis, Union City, California). Bats’ body temperatures were obtained via implanted transponder (BMDS IPTT-300) and weights were taken on days −8/7, 2, 4, 6, 8, 10, 12, 14, 17, 21, 24, and 28. Swabs were collected in 1 mL DMEM2. Bats were observed daily for clinical signs of disease. Necropsies and tissue sampling were performed according to IBC-approved SOPs.

### In vitro infections

To assess filovirus replication and host gene expression kinetics, AjKi_RML2 and RoNi cells were plated at 250,000 cells/mL in 10DMEM/F-12 or DMEM10, respectively, in 24-well plates and incubated overnight at 37°C, 5% CO_2_. The next day, medium was removed, and cells were inoculated with either EBOV-Mayinga or MARV-Ozolin at multiplicity of infection (MOI) 0.1 for 1 hour at 37°C, 5% CO_2_ with rocking every 15 minutes. Cells were then washed three times with 1X Dulbecco′s phosphate buffered saline, no calcium, no magnesium (DPBS) and 0.5 mL of fresh medium, DMEM/F-12 with 2% FBS, 1X NEAA, and 50 U/mL penicillin, and 50 μg/mL streptomycin (2DMEM/F-12) for AjKi_RML2 and DMEM2 for RoNi, was added. A sample of the medium was collected at the one-hour time point, and the volume was replaced with fresh medium. Every 24 hours for five days, 500 μL of supernatant was collected for titration and the cells were lysed in 600 μL of RLT (QIAGEN) for RNA extraction or 4% sodium dodecyl sulfate (SDS) buffer with 10% β-mercaptoethanol for western blotting. Protein samples were boiled at 100°C for 10 minutes. Both supernatant and RLT samples were stored at −80°C until processing. The same protocol was used with EBOV-Kikwit and EBOV-Makona C05 and MARV-Angola and MARV-Musoke in additional JFB and ERB cell lines at 72 and 120 HPI.

To evaluate antagonism of IFN-I signaling, AjKi_RML2 and Huh-7 cells were plated at 250,000 cells/mL in 6-well plates in 2.0 mL of the appropriate medium and incubated overnight at 37°C, 5% CO_2_. The next day, medium was removed, and cells were inoculated with MOI 1.5 of virus (EBOV-Mayinga or MARV-Angola) for 1 hour at 37°C, 5% CO_2_ with rocking every 15 minutes. After incubation, the inoculum was removed and 2.0 mL of fresh medium was added and cells were returned to the incubator for 24 hours. Cells were stimulated with 100 ng/mL of Chiroptera IFNβ for 30 minutes or mock treated with equal volume of medium. Cells were trypsinized for 10 minutes with TrypLE Express (Gibco) and spun down after addition of an equal volume of complete medium. After washing the cell pellet once with 1X DPBS, the cytoplasm and nucleus were fractionated using NE-PER™ Nuclear and Cytoplasmic Extraction Reagents (Thermo Scientific, Cat # 78833) according to the manufacturer’s instruction. The nuclear and cytoplasmic fractions were inactivated according to SOP and western blot analysis was performed as described below.

### IFNβ expression and purification

A bat *Ifnb1* consensus sequence was designed using the *Ifnb1* gene sequence from all bat genomes available in the NCBI database as of November 2022. To prevent bias from the number of species represented in different bat families, we first generated a consensus of each bat family and generated the Chiroptera consensus from the bat family consensus sequences. An IL-2 secretion signal, His-tag, and TEV protease signal were added upstream of the consensus *Ifnb1* (Chiroptera *Ifnb1*) open reading frame and the synthesized sequence was inserted into the pcDNA3.4 vector (GenScript).

tsA201 cells derived from HEK293 (Millipore Sigma, cat# 96121229-1VL), from which *Mycoplasma* was eradicated at ECACC, and the identity of tsA201 and 293 has been confirmed by STR (short tandem repeat) profiling. Cells were adapted to grow in suspension by using Gibco® FreeStyle™ 293 Expression Medium plus 2 mM glutamine in a shaker flask. The low passage (P2 or P3) of the suspension culture was collected, and cell stock was made with the addition of 7.5% DMSO and then stored in liquid nitrogen. For each expression, one frozen vial with 1×10^7^ tsA201 cells was inoculated with 40 mL of the above-mentioned medium in a 250 mL flat-bottom cell culture shaker flask. Cell density was measured after cells grew for three days in an Infors Multitron incubator at 130 rpm with 8% CO_2_ and 75% humidity, using NucleoCounter® NC-100™. Cells were then passed into a large flask (2.8 L or 5 L Optimum Growth Flask) with the addition of the above-mentioned medium to reach a final cell density of 1.3 x 10^5^/mL. The total volume of culture was no more than 1.0 L for a 2.8 L flask and no more than 2.1 L for a 5 L flask. Transfection was started when, cells reached a density of 2 × 106/mL, typically after 3 days. For one liter of culture, 1 mg of pcDNA3.4-Chiroptera-*Ifnb1* plasmid (filtered with a 0.22 µm filter) was added to 50 mL of prewarmed Hybridoma-SFM (ThemoFisher, 12045076). Four milliliters of PEI MAX® (Transfection Grade Linear Polyethylenimine Hydrochloride, MW 40,000, Polysciences, 24765-1) at 1 mg/mL was added, then gently mixed to homogenize. After incubation at room temperature for 12–15 minutes, the mixture was added to the culture, and then the flask was returned to the incubator. After five days, the culture was centrifuged at 1,000 g for 15 minutes using a Sorvall RC3B plus centrifuge. The supernatant was stored at −80°C for future purification.

The supernatant was filtered through a 0.22 µm filter, then mixed with 5 mL of prewashed HisPur™ Ni-NTA Resin (ThermoFisher Scientific) and rocked overnight at 4°C. The following day, the supernatant was drained, and the resin was packed into a 5 mL column. AKTA FPLC with Unicorn software (GE Healthcare) was used to wash the column with sodium phosphate buffer, pH 7.0, and 200 mM NaCl at 2 mL/min. The protein was eluted with a linear gradient up to 400 mM imidazole over 30 CV. The IFN-β-containing fractions were pooled (∼28 mL) and injected into a 30 mL Slide-A-Lyze cassette with 10K MWCO (ThermoFisher Scientific). The cassette was then dialyzed against 1.8 L of 50 mM sodium phosphate buffer, pH 7, three times. After dialysis, the protein concentration was measured using a NanoDrop Microvolume Spectrophotometer (ThermoFisher Scientific). IFN-β was then cleaved by TEV protease to remove the N-terminal His tag. The enzymatic cleavage was completed with 1 mg of TEV per 5 mg of IFN-β and incubated at 4°C overnight with rocking. The digested mixture was then loaded onto a pre-equilibrated 5 mL Ni-NTA Superflow column (Qiagen) to remove the cleaved His-tag, remaining uncleaved protein, as well as additional impurities, with a flow rate of 1 mL/min. The flowthrough fractions were loaded onto a pre-equilibrated HiTrap Capto Q anion exchange column (Cytiva) to separate IFN-β aggregates from monomer with a flow rate of 2 mL/min. The running buffer was 50 mM sodium phosphate, pH 7, and the gradient was 5 mM/min for NaCl from 0 to 500 mM. IFN-β monomer-containing fractions around 200 mM NaCl were immediately collected and analyzed with SDS-PAGE.

### IFN-I pathway inhibition

AjKi_RML2 cells were treated with 5-fold serial dilution of IFN-I induction (BAY-985), signaling (deucravacitinib, fludarabine, or itacitinib) inhibitors or dimethyl sulfoxide (DMSO) vehicle control for 24 hours prior to stimulation with 500 ng/mL high molecular weight poly(I:C) (Invitrogen) for 18 hours (BAY-985) or 100 ng/mL recombinant consensus bat IFN-β (IFN-I signaling inhibitors) for 30 minutes. Protein was collected after stimulation and the effect of blocking IFN-I signaling on filovirus replication was assessed after infection of cells with MOI 0.1 of EBOV-Mayinga or MARV-Ozolin, as described above, then added itacitinib in 10-fold serial dilution (0.25-25nM) or equal volume of DMSO. The medium was replaced with fresh inhibitor every 24 hours before collection of supernatants for titration at 72 hours post-infection.

### IFN-β sensitivity

To assess the sensitivity of EBOV and MARV to IFN-β, AjKi_RML2 or RoNi cells were treated with 10-fold serial dilutions of Chiroptera IFN-β (0.1-100 ng/mL) for 24 hours prior to infection with MOI 0.1 of EBOV-Mayinga or MARV-Ozolin. Uninfected cells were lysed in RLT 24 hours after IFN-β stimulation to measure the induction of interferon stimulated genes at the time of infection. Supernatants were collected 48 hours after infection and titrated on Vero E6 cells as described above.

### Western blot analysis

Inactivated cell lysates collected in SDS buffer were run on 4-12% Bis-Tris NuPAGE gels (Invitrogen) at 150V for 1 hour then transferred onto methanol-activated PVDF membrane (BioRad). Following a 1 hour block in 5% powdered milk, the membranes were washed in buffer (1X tris-HCL with 0.1% tween 20) three times. Membranes were probed with primary antibody overnight rocking at 4°C. The next day, blots were washed three times, incubated with an HRP-conjugated secondary antibody for 1 hour rocking at room temperature, and washed three times before development. Blots were incubated with a 1:1 ratio of peroxidase and enhancer reagents Clarity Western ECL (BioRad) or SuperSignal West Femto (Thermo Scientific) and developed on an iBright imaging system (Thermo Fisher Scientific). To calculate relative expression, area under the curve was determined for each band using Fiji [35]. Primary antibodies used: pSTAT1 – Y701 (Cell Signaling Technology, 9167S), pSTAT2 – Y690 (Cell Signaling Technology, 88410S), total STAT1 (Cell Signaling Technology, 14994S), total STAT2 (Cell Signaling Technology, 72604S), RIG-I (Kerafast, 1C3), Lamin A/C (Cell Signaling Technology, 4777S), β-tubulin (Sigma Aldrich, T8328), EBOV VP24 (Sino Biological 40454-T46), EBOV VP40 (GeneTex, GTX134034), MARV VP40 (The Native Antigen Company, MAV12450-100). Secondary antibodies used: Donkey-anti-rabbit (GE Healthcare, NA934) and Sheep-anti-mouse (GE Healthcare, NA931). Area under the curve (AUC) for western blot bands was measured with Fiji [35]. Normalized expression was determined with phosphorylated protein with the following formulas: (AUC pSTAT1/AUC β-tubulin or Lamin A/C) or (AUC VP40/ AUC β-tubulin). To calculate relative expression, the normalized expression of the sample is divided by the normalized expression of the mock/non-infected value.

### Pseudoparticle assay

Pseudoparticles were generated using VSVs containing the GFP gene instead of the receptor-binding VSV G protein gene (VSVΔG) pseudotyped with EBOV-Mayinga GP or Marburg-Musoke GP as described previously [36]. Infectious units (IUs) of these pseudotyped VSVs were determined on Vero E6, AjKi_RML1, AjKi_RML2, and RoNi cells using methods previously described [37]. The normalized entry ratio was calculated for each cell line using the following formula: ((IU EBOV/IU MARV) for cell line of interest)/((IU EBOV/IU MARV) for Vero E6).

### Recombinant VSV infections

Stocks of replicating recombinant VSV (rVSV)-GFP encoding GPs from wild-type VSV, EBOV-Mayinga, or MARV-Ozolin were titrated on either Vero E6 cells or AjKi_RML2 to determine differential TCID_50_. To evaluate replication kinetics, Vero E6 or AjKi_RML2 cells were infected at MOI 0.005 with VSV-MARVozolin-GFP and - EBOVmayinga-GFP for 1 hour at 37°C, 5% CO_2_ with constant rocking. Cells were washed three times with 1 mL of DPBS before the addition of fresh medium. The supernatants were titrated on Vero E6 cells.

### RNA extractions and RT-qPCR

Swabs and blood were collected as described above; 140 µL was used for RNA extraction using the QIAamp Viral RNA Kit (Qiagen) according to the manufacturer’s instructions with an elution volume of 60 µL. For tissues and cells, RNA was isolated using the RNeasy Mini kit (Qiagen) according to the manufacturer’s instructions and eluted in 50 µL. Viral RNA was detected by RT-qPCR (**Supplementary Table 1**). RNA was tested with TaqMan™ Fast Virus One-Step Master Mix (Applied Biosystems) using QuantStudio 6 or 3 Flex Real-Time PCR System (Applied Biosystems). EBOV or MARV standards with known copy numbers were used to construct a standard curve and calculate copy numbers/mL or copy numbers/g.

RLT lysates from cell monolayers were transferred to 70% ethanol prior to extraction of RNA using the RNeasy extraction kit (Qiagen). Cellular RNA was used to measure viral RNA and host genes using RT-qPCR (**Supplementary Table 1**). Following extraction, 17 µL of RNA was treated with TURBO DNase (Invitrogen) according to the manufacturer’s instructions. After DNase treatment, the RNA was diluted 1:5 in molecular grade water (Invitrogen). DNase-treated RNA (5 µL) was used for each host gene assessed. Fold change was calculated for host genes by dividing the sample relative gene expression by the average relative gene expression of the healthy controls or mock-treated cells (2^-ΔΔCt^).

### Virus titration

Infectious virus in tissue, swab, and supernatant samples was evaluated in Vero E6 cells. Tissue samples were weighed, then homogenized in 1 mL of DMEM2. Vero E6 cells were inoculated with ten-fold serial dilutions of homogenate, swabs, blood, or supernatant incubated for 1 hour at 37°C. For tissue homogenates and blood, the first two dilutions of each sample replicate were washed twice with DMEM2. For swab and supernatant samples, cells were inoculated with ten-fold serial dilutions and no wash was performed. On days 10-14, cells were scored for cytopathic effect (CPE). Titers in TCID_50_/mL were calculated by the Spearman-Karber method.

### Serology

Maxisorp plates (Nunc) were coated with 50 ng of EBOV-Mayinga [38] or MARV-Angola (IBT Bioservices, cat no. 0506-015) GP per well. Plates were incubated overnight at 4°C. Plates were blocked with casein in phosphate buffered saline (PBS) (ThermoFisher) for 1 hour at room temperature. Sera were diluted 1:100 in blocking buffer and samples (triplicate) were incubated for 1 hour at room temperature. Secondary horseradish peroxidase (HRP)-conjugated recombinant protein-A/G (Invitrogen, lot number WH 324034) diluted 1:10,000 was used for detection and visualized with KPL TMB two-component peroxidase substrate kit (SeraCare, 5120-0047). The reaction was stopped with KPL stop solution (SeraCare) and plates were read at 450 nm. Plates were washed three times with PBS-T (0.1% Tween) between each step. Arbitrary enzyme-linked immunosorbent assay (ELISA) units were calculated based on the standard curve. All samples diluted at 1:100 with an optical density value of <0.250 were given a value of 1.

### Virus neutralization

Heat-inactivated, irradiated sera were two-fold serially diluted in DMEM2, and 100 TCID_50_ of EBOV-Mayinga or MARV-Ozolin was added. After 1 hour of incubation at 37°C and 5% CO_2_, the virus:serum mixture was added to Vero E6 cells. CPE was scored after 11-14 days incubated at 37°C and 5% CO_2_. Viral neutralization was determined based on the dilution factor non-CPE wells were observed.

### Flow cytometry

Bat cells were isolated from spleens by generating a single cell suspension using a 100-µm filter and red blood cells were removed using RBC lysis buffer (Invitrogen). Splenocytes were then cryopreserved in freezing medium (10% DMSO and 90% FBS) and stored at −80°C. The cryopreserved bat splenocytes were rapid thawed in RPMI-1640/10% FBS and washed twice. To test for presence of EBOV GP-specific B cells, Alexa Fluor 568 was conjugated to recombinant EBOV GP receptor binding domain (Sino Biological, 40304-V08H) using an Invitrogen Alexa Fluor protein labeling kit (Invitrogen, A10238). They were first stained with ViaKrome 808 Fixable Viability Dye (Beckman Coulter) for 30 minutes at room temperature (RT) and then surface stained with EBOV-GP AF568 for 30 minutes at RT. Samples were then fixed and permeabilized using a Cytofix/Cytoperm kit (BD Biosciences). The antibodies used for intracellular staining were: Allophycocyanin-anti-CD79a (HM57, BD Biosciences 752115) and Fluorescein isothiocyanate-anti-CD3ε (CD3-12, BioRad MCA1477F). Samples were analyzed on a 6-laser Cytoflex LX (Beckman Coulter) and flow cytometry data were analyzed using FlowJo v.10.9 (BD Life Sciences). Live lymphocytes were gated using a FSC-A vs Live-Dead gate, followed by a FSC-A and FSC-H gate to exclude doublets and then by a standard FSC-A vs SSC-A lymphocyte gate (**Supplementary Figure 1**).

### B cell receptor sequencing

Splenic mRNA was harvested from healthy control JFBs and from EBOV infected bats at 3-, 7-, and 28 DPI. The mRNA was inactivated and stored in Trizol according to institutional SOP. After addition of chloroform to Trizol, RNA was extracted with Phasemaker tubes (Invitrogen). For BCR library preparation, we followed previous protocols for JFB BCR sequencing (Crowley, 2024). We prepared BCR libraries with the NEBNext® Ultra II DNA Library Prep Kit with Sample Purification Beads (cat# E7103S) and NEBNext® Multiplex Oligos for Illumina® (Index Primers Set 1, cat# E7335S). Samples were cleaned using NebNext Ultra II beads and enriched for libraries with a size range of 500 to 700 base pairs amplicons, according to manufacturers instructions [39]. All libraries were sequenced using the Illumina MiSeq 2×300 chemistry platform.

Sequencing reads were cleaned using Immcantation version 4.3.0. Sequences were filtered out if they had an average FASTQ score below 30. Sequences with shared unique molecular identifiers were used to generate consensus sequences. Consensus sequences with only one contributing read were removed from the dataset. The *Mus musculus* germline was used to classify V, D, and J mRNA sequences into putative germline genes using IgBLAST (version 1.18.0[40]. The mouse germline was used because the V, D, and J germline genes have not been fully annotated for the JFB.

Consensus sequences were binned into clonal lineages with a Hamming distance cutoff of 0.09. The frequency of clones and V_H_ and J_H_ gene segments were assessed using Alakazam (version 1.2.1). Clonal frequencies were normalized with an ordered quantile normalization transformation using the R library BestNormalize (version 1.9.1). For each bat, a subsample of 300 clones was randomly selected. These subsamples were then analyzed using a Bayesian hierarchical model implemented in Rstan (version 2.32.5). The model estimated the average normalized clonal frequency for each time point based on the data from the bats euthanized at those respective time points. The model pooled information across individuals to estimate a population-level average clonal frequency for each time point, while accounting for individual variation.

### Histopathology

Necropsies and tissue sampling were performed according to IBC-approved SOPs. Tissues were collected and fixed for a minimum of 7 days in 10% neutral buffered formalin with two formalin changes. Tissues were placed in cassettes and processed with a Sakura VIP-6 Tissue Tek on a 12-hour automated schedule using a graded series of ethanol, xylene, and PureAffin. Prior to staining, embedded tissues were sectioned at 5 µm and dried overnight at 42°C. Histopathologic analysis was performed by a blinded, board-certified veterinary pathologist.

### Next-generation sequencing of mRNA

RNA from livers of negative control or 3 DPI EBOV- or MARV-infected bats was used for analysis. Two hundred nanograms of RNA was used as input for poly-A pull-out and next-generation sequencing (NGS) library preparation following the Illumina Stranded mRNA prep workflow (Illumina). The NGS libraries were prepared, amplified for 15 cycles, purified with AMPureXP beads (Beckman Coulter), assessed on a TapeStation 4200 (Agilent Technologies), and quantified using the Kapa Library Quantification Kit (Illumina). Amplified libraries were normalized, pooled at equal 2 nM concentrations, and sequenced as 2 × 75 base reads on the NextSeq instrument using three high output chemistry kits (Illumina). Raw fastq reads were trimmed of Illumina adapter sequences using cutadapt (version 1.12), and then trimmed and filtered for quality using the FASTX-Toolkit (Hannon Lab). Remaining reads were aligned to the Jamaican fruit bat genome assembly version (GCF_021234435.1) using Hisat2 [35]. Reads mapping to genes were counted using htseq-count [36]. Differential expression analysis was performed using the Bioconductor package DESeq2 [37] and data were further analyzed and plotted using ggplot2 (V3.4.0) as part of the tidyverse package (V1.3.2) [38]. Pathway analysis was performed using Ingenuity Pathway Analysis (Qiagen), and gene clustering was performed using Partek Genomics Suite (Partek Inc.).

### Statistical Analysis

Significance tests were performed as indicated where appropriate for the data using GraphPad Prism 10.2.0. Unless stated otherwise, statistical significance levels were determined as follows: no symbol = p > 0.05; * = p ≤ 0.05; ** = p ≤ 0.01; *** = p ≤ 0.001; **** = p ≤ 0.0001. The statistical test used is specified where appropriate.

### Data availability statement

All data are available on request from the corresponding authors or will be uploaded to FigShare. All material requests should be sent to V.J.M..

## Results

### Infection of JFBs with EBOV or MARV does not cause overt clinical disease

Bats were inoculated with EBOV (n=12) or MARV (n=12) via the oral, intranasal, and subcutaneous routes. At 3- and 7-DPI, four EBOV- and four MARV-infected bats were euthanized, and the remaining animals were monitored through 28 DPI (**Figure 1A**). The EBOV-infected bats developed mild hypothermia with a statistically significant decrease in body temperature at 4-, 8-, and 10-DPI relative to baseline temperatures (**Figure 1B**). All EBOV-infected bats returned to baseline body temperatures by 28 DPI. There were no statistically significant changes in body weight compared to baseline in EBOV-infected bats (**Figure 1C**). The MARV-infected bats did not have significant changes in body temperature (**Figure 1D**) or weight (**Figure 1E**). MARV05 was pregnant and gained weight throughout the study.

**Figure 1.**
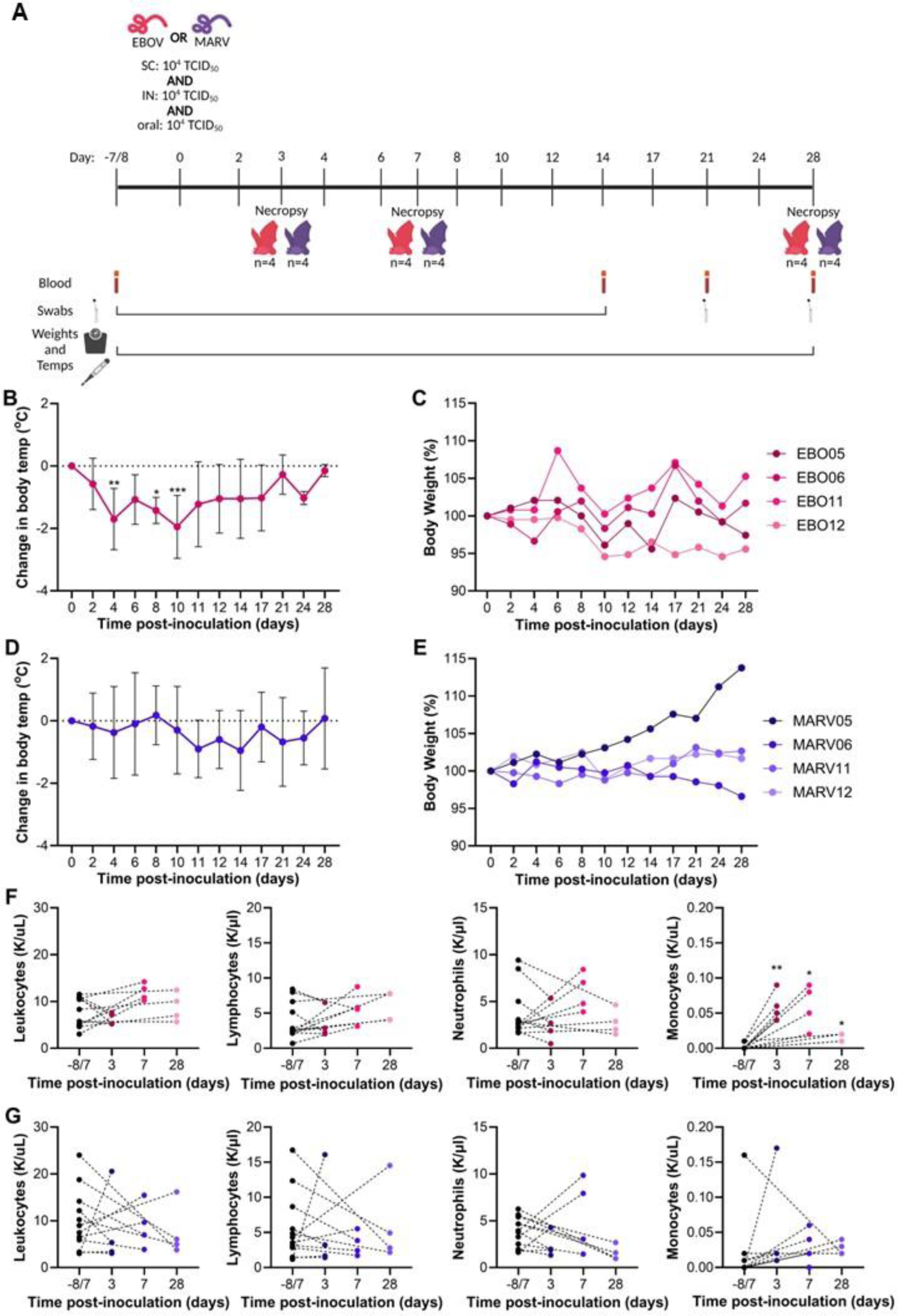
Infection of JFB with EBOV or MARV does not cause overt clinical disease. (A) Depiction of challenge study of Jamaican fruit bats with EBOV-Mayinga (magenta) or MARV-Ozolin (purple) via subcutaneous (SC), intranasal (IN), and oral routes. Image created using Biorender. (B) Change in body temperature of bats (n=4) infected with EBOV followed through day 28. Two-way ANOVA with Dunnett’s multiple test comparison. (C) Percent body weight of bats (n=4) infected with EBOV followed through day 28. Two-way ANOVA with Dunnett’s multiple test comparison. (D) Change in body temperature of bats (n=4) infected with MARV followed through day 28. Two-way ANOVA with Dunnett’s multiple test comparison. (E) Percent body weight of bats (n=4) infected with MARV followed through day 28. Two-way ANOVA with Dunnett’s multiple test comparison. (F) Number of circulating white blood cells (WBC), lymphocytes, neutrophils, and monocytes in EBOV-infected bats at baseline and at necropsies. A mixed-effects model with matching and Dunnett’s multiple comparison test was applied to evaluate change in each cell population at necropsy compared to the matched baseline value. (G) Number of circulating WBC, lymphocytes, neutrophils, and monocytes in MARV challenged bats at baseline and at necropsies. A mixed-effects model with matching and Dunnett’s multiple comparison test was applied to evaluate change in each cell population at necropsies compared to the matched baseline value.

Since lymphopenia is a hallmark of filovirus disease in pathogenic hosts [41, 42], we collected whole blood at necropsy for complete blood cell counts. There were no significant changes in total white blood cells (WBC), lymphocytes, or neutrophils relative to baseline in either EBOV- or MARV-infected bats (**Figure 1F, G**). In EBOV-infected bats, the number of monocytes increased significantly at 3- and 7-DPI (**Figure 1F**). The number of circulating monocytes did not change after MARV infection (**Figure 1G**). We did not observe any other statistically significant changes in hematology (**Supplementary Table 2**) or serum chemistries (**Supplementary Table 3**).

### JFBs shed infectious EBOV orally

Prior to infection, every two days through 14 DPI, 21 DPI, and at necropsies we collected oropharyngeal and rectal swabs to monitor viral shedding (**Figure 1A**). EBOV RNA was detected in oropharyngeal swabs 2-10 DPI, peaking at 6 DPI (**Figure 2A**), and rectal swabs 2-6 DPI (**Figure 2B**). MARV RNA was not detectable in any oropharyngeal or rectal swabs (**Figure 2C, D**). Infectious EBOV was present in oropharyngeal swabs, up to 3.2 log_10_ TCID_50_/mL (**Figure 2E**), but not in rectal swabs (**Figure 2F**). No infectious MARV was detected in 6 DPI oropharyngeal swabs (**Figure 2G**).

**Figure 2.**
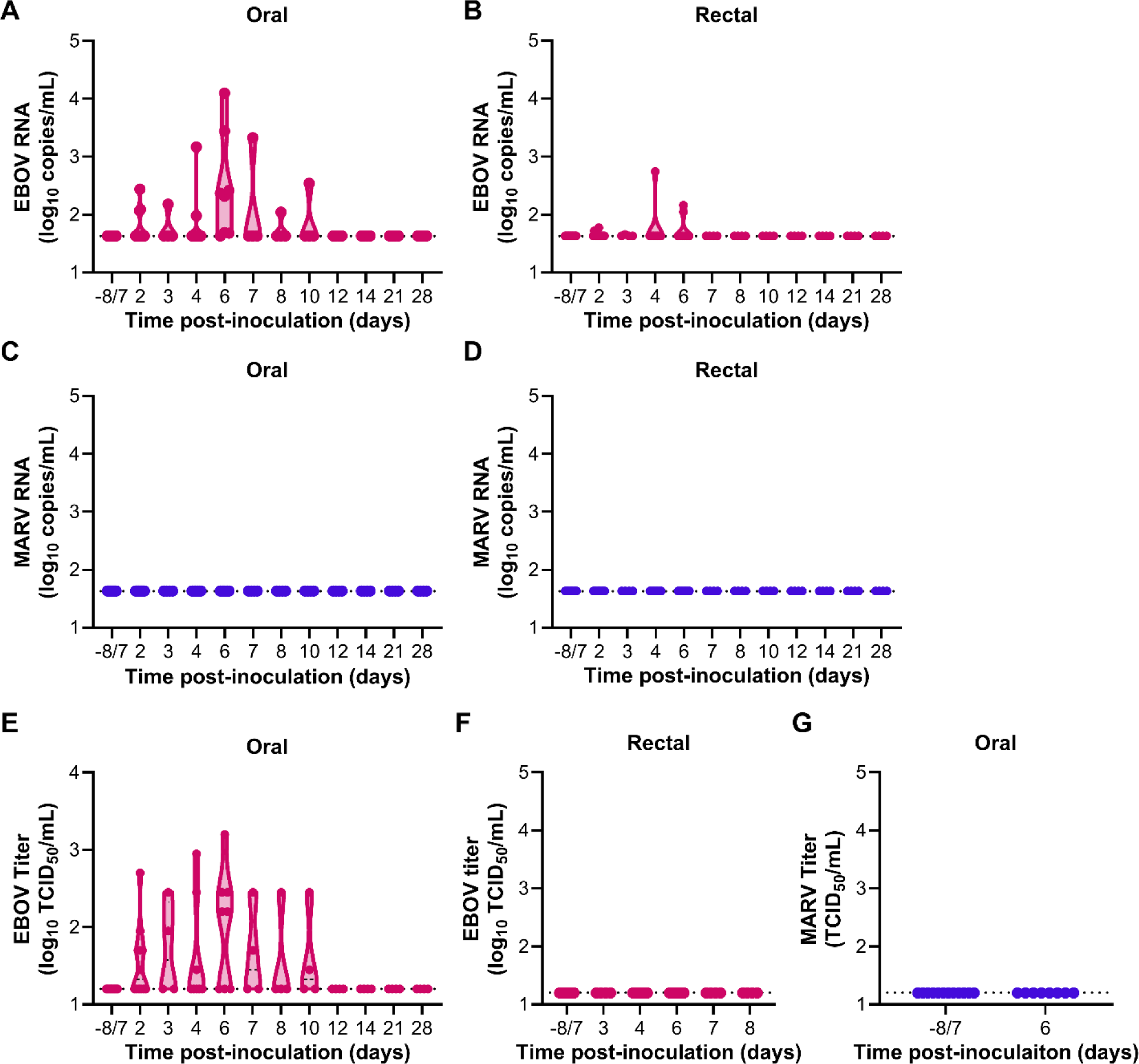
Infected JFB shed infectious EBOV orally. (A) RT-qPCR for EBOV RNA in the oral swabs. Limit of detection 1.63log_10_ copies/mL. (B) RT-qPCR for EBOV RNA in the rectal swabs. Limit of detection 1.63log_10_ copies/mL. (C) RT-qPCR for MARV RNA in the oral swabs. Limit of detection 1.63log_10_ copies/mL. (D) RT-qPCR for MARV RNA in the oral swabs. Limit of detection 1.63log_10_ copies/mL. (E) Infectious titer of EBOV in the oral swabs. Limit of detection 1.2log_10_ TCID_50_/mL. (F) Infectious titer of EBOV in the rectal swabs in span of days where viral RNA was detected in swabs. Limit of detection 1.2log_10_ TCID_50_/mL. (G) Infectious titer of MARV in the oral swabs at baseline and on day 6 post-challenge. Limit of detection 1.2log_10_ TCID_50_/mL.

### EBOV, but not MARV infection is disseminated in JFBs

Tissues collected on the sequential necropsy days were analyzed for the presence of viral RNA, infectious virus, and viral antigen. EBOV RNA was detected in all tissues evaluated, skin at the inoculation site (SIS) (8/8), salivary gland (5/8), lung (7/8), liver (8/8), spleen (7/8), kidney (5/8), and reproductive organs (3/8) (ovary for female, testis for male), for at least one bat at 3 and 7 DPI necropsies (**Figure 3A**). The highest levels of EBOV RNA were detected in the SIS (5.5-7.5 log_10_ copies/g) at 3 and 7 DPI (**Figure 3A**). EBOV RNA remained detectable in the spleens (4.5-5.9 log_10_ copies/g) of all four bats and one SIS sample (2.9 log_10_ copies/g) at 28 DPI while the other tissues were negative (**Figure 3A**). Titration of the tissue samples largely reflected the RT-qPCR data with infectious virus present in at least one sample for all tissues tested (**Figure 3B**). We were unable to evaluate the presence of infectious virus in spleen samples collected at 28 DPI due to insufficient sample. Minimal MARV RNA was detected in the SIS (8/8, 2.7-4.7 log_10_ copies/g), liver (1/8, 2.7 log_10_ copies/g), and spleen (2/8, 3.5-4.6 log_10_ copies/g) at 3 and 7 DPI, and all other samples were negative (**Figure 3C**). Minimal infectious MARV was detected in SIS and liver samples collected at 3- and 7 DPI (**Figure 3D**).

**Figure 3.**
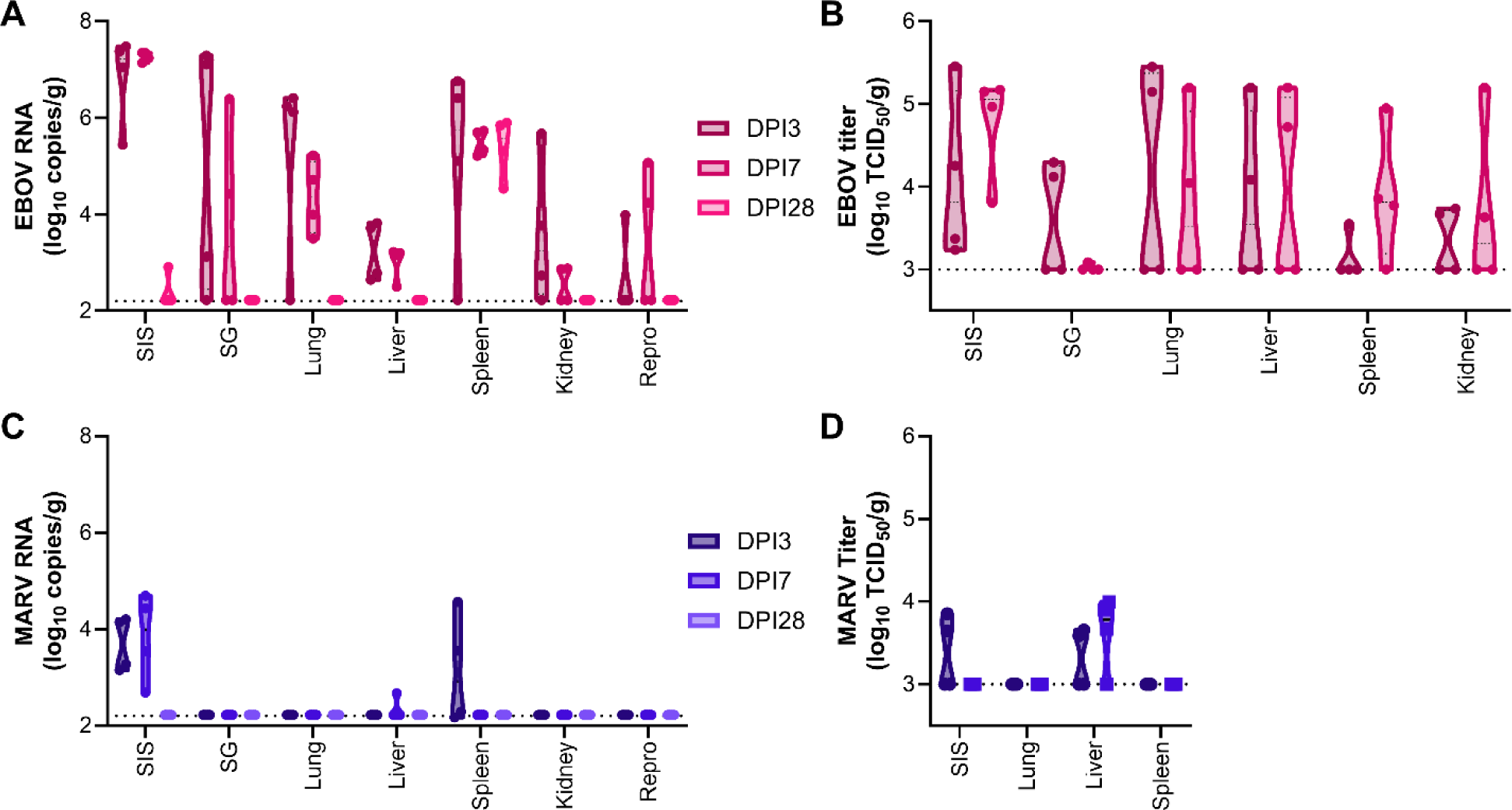
EBOV but not MARV infection is disseminated in JFB. (A) RT-qPCR for EBOV in tissue samples collected at 3-, 7-, and 28 days post-infection (DPI) (n=4). Limit of detection 2.2log_10_ copies/g. (B) Infectious titer of EBOV in tissues collected at D3, 7, and 28 (n=4). Limit of detection 3.0log_10_ copies/g. (C) RT-qPCR for MARV in tissue samples collected at necropsy day (DPI) 3, 7, and 28 (n=4). Limit of detection 2.2log_10_ copies/g. (D) Infectious titer of MARV in tissues collected at D3, 7, and 28 (n=4). Limit of detection 3.0log_10_ copies/g.

Injection site skin, salivary gland, cervical lymph node, soft palate/tonsil, lung, heart, thymus, liver, spleen, kidney, adrenal gland, urinary bladder, gonad, reproductive tract, gastrointestinal tract and eye were evaluated by histopathologic analysis for evidence of viral induced inflammation (**Supplementary Figure 2**). In EBOV-infected bats, sections of SIS revealed granulomatous inflammation in the subcutis. Subcuticular inflammation was most pronounced in bats 417, 419, 423 and 424, reaching moderate inflammation. When present for evaluation, neutrophilic lymphadenitis was observed in cervical lymph nodes ranging from minimal to moderate severity. Sporadic splenic changes were observed at all timepoints in EBOV-infected bats including minimal to moderate splenic lymphoid depletion (n=8) and minimal to mild lymphoid follicular necrosis with neutrophilic influx (n=5). In MARV-infected bats, minimal to mild subcutaneous perivascular cuffing, pyogranulomatous panniculitis, and focal myocyte regeneration were observed sporadically at all evaluated timepoints. No significant histopathologic lesions were observed in evaluated sections of tonsil/palate, salivary gland, kidney, adrenal gland, lung, heart, thymus, liver, urinary bladder, gastrointestinal tract or eye from either EBOV- or MARV-infected JFBs.

### EBOV infection induces robust innate antiviral and humoral responses

Host gene expression analysis using RT-qPCR was performed on SIS, lung, liver, and spleen samples collected at 3- and 7 DPI necropsies for both interferon stimulated genes (ISGs) and inflammatory cytokines. In the SIS, expression of ISGs, *Isg15*, *Ifit1*, *Mx1*, *Oas1*, and *Ddx58*, was induced significantly in EBOV-infected bats at 3 DPI and remained marginally elevated at 7 DPI (**Figure 4A**). EBOV infection significantly induced *ll6* expression in the SIS at 7 DPI compared to healthy control bats (**Figure 4B**). In MARV-infected bats, *Ifit1* at 3 DPI was induced and *Il1b* mRNA expression was inhibited at 3 and 7 DPI compared to the healthy controls (**Figure 4A, B**). At 3 DPI with EBOV, all ISG mRNAs, except RIG-I, were significantly induced in the liver (**Figure 4C**). Expression of *Il1b* (7 DPI) and *Tnfa* (3 and 7 DPI) were significantly induced in the liver of EBOV-infected bats (**Figure 4D**). *Ifit1* (3 DPI) and *Tnfa* (3- and 7 DPI) were elevated significantly in the liver of MARV-infected bats (**Figure 4C, D**). In the 3 DPI spleens, all ISG mRNAs were induced in EBOV-infected bats and *Ifit1* was induced in MARV-infected bats (**Figure 4E**). *Tnfa* was significantly induced in the spleen of EBOV- and MARV-infected bats 7 DPI, and *Il6* mRNA was elevated significantly at 3 and 7 DPI after EBOV infection (**Figure 4F**). In the lung *Isg15*, *Ifit1*, and *Mx1* were the only ISGs with significantly increased transcription in EBOV-infected bats, and ISG expression in MARV-infected bats did not differ from healthy controls (**Supplementary Figure 3A**). Proinflammatory cytokine genes were not differentially expressed in either EBOV- or MARV-infected bats at either timepoint (**Supplementary Figure 3B**). Of note, the magnitude of induction for ISGs was higher than for pro-inflammatory cytokines in all tissues.

**Figure 4.**
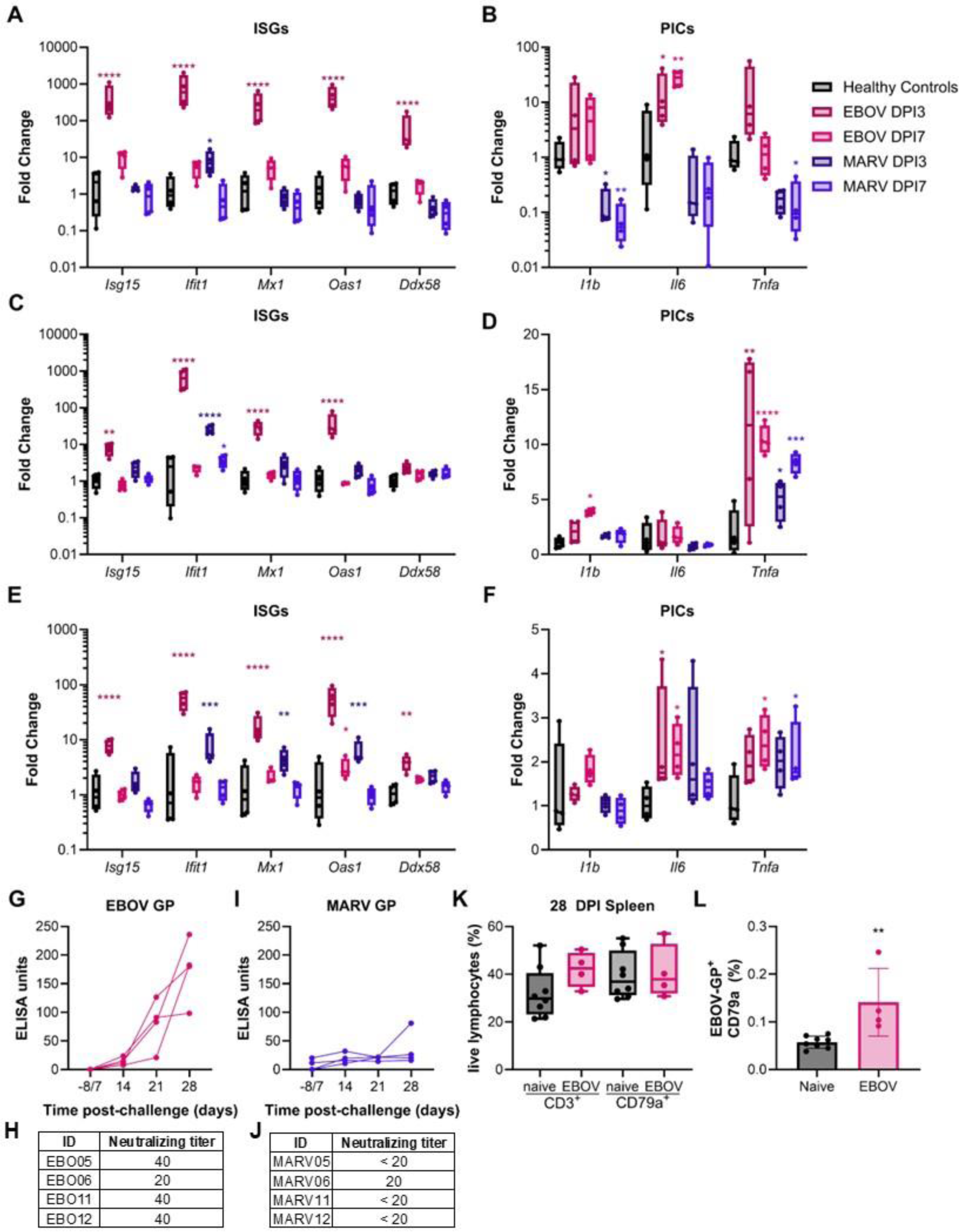
EBOV infection induces robust innate antiviral and adaptive immune responses. (A) RT-qPCR of interferon stimulated gene (ISG) mRNA in the skin at inoculation site collected from healthy control bats (n=4) and at day (D) 3 or 7 necropsy of EBOV or MARV-infected bats (n=4). (B) RT-qPCR of pro-inflammatory cytokine gene (PIC) mRNA in the skin at inoculation site collected from healthy control bats (n=4) and at day (D) 3 or 7 necropsy of EBOV or MARV virus challenged bats (n=4). (C) RT-qPCR of ISG mRNA in livers collected from healthy control bats (n=4) and at day (D) 3 or 7 necropsy of EBOV or MARV virus challenged bats (n=4). (D) RT-qPCR of PIC mRNA in livers collected from healthy control bats (n=4) and at day (D) 3 or 7 necropsy of EBOV or MARV virus challenged bats (n=4). (E) RT-qPCR of ISG mRNA in spleens collected from healthy control bats (n=4) and at day (D) 3 or 7 necropsy of EBOV or MARV virus challenged bats (n=4). (F) RT-qPCR of PIC mRNA in spleens collected from healthy control bats (n=4) and at day (D) 3 or 7 necropsy of EBOV or MARV virus challenged bats (n=4). (A-F) ΔC_T_ values were normalized to the average ΔC_T_ value of the healthy control bats to calculate ΔΔC_T_. Fold change of each gene at necropsy day was compared to the healthy controls using a two-way ANOVA with Dunnett’s multiple test comparison. (G) Serum binding antibodies and (H) neutralizing titers were determined via ELISA assay with a standard curve of EBOV glycoprotein (GP) or neutralization assay, respectively. Neutralizing titer is presented as the lowest dilution that protected 50% of wells in a TCID_50_ assay. (I) Serum binding antibodies and (J) neutralizing titers were determined via ELISA assay with a standard curve of MARV GP or neutralization assay, respectively. Neutralizing titer is presented as the lowest dilution that protected 50% of wells in a TCID_50_ assay. (K) Cryopreserved splenocytes from naïve controls (n=8) and D28 EBOV (n=4) infected bats were analyzed for intracellular expression of CD3ε and CD79a to identify the proportion of T cells and B cells from a live lymphocytes gate. (L) CD79a+ B cells from cryopreserved splenocytes were analyzed for interaction with Alexa-fluor 598 conjugated recombinant EBOV-GP receptor binding domain. The percentage of live CD79a^+^ B cells with a positive EBOV-GP staining signal after subtracting the fluorescence minus one control signal.

We performed transcriptomic analysis on 3 DPI liver samples because we observed viral RNA and infectious virus for both EBOV- and MARV-infected bats. Comparing EBOV-infected bats with healthy controls, we found 241 upregulated and 460 downregulated genes; comparing MARV-infected bats with healthy controls, we found 230 upregulated and 516 downregulated genes (significance level = 0.05, foldchange >2). The Gene Ontology enrichment analysis revealed that nearly all the top ten pathways enriched in EBOV-infected bats were also enriched in those infected with MARV and are involved in the regulation of the immune response (**Supplementary Table 4**). When comparing EBOV- and MARV-infected bats, we observed 98 upregulated and 146 downregulated genes in EBOV-relative to MARV-infected bats (significance level = 0.05, foldchange >2). The pathways differentially regulated between the two filoviruses are involved in immune regulation (**Supplementary Table 5**). Interestingly, some genes involved in antigen processing and presentation are induced in EBOV-but not MARV-infected bats.

GP-specific antibodies were measured using ELISA on serum collected from the bats monitored through 28 DPI. All four of the EBOV-infected bats seroconverted with EBOV GP-specific binding antibodies steadily increasing after infection (**Figure 4G**). The titers of EBOV neutralizing antibodies at 28 DPI were 20 for EBO420 and 40 for the other bats (**Figure 4H**). Only one of the four MARV-infected bats, MARV80, seroconverted (**Figure 4I**). MARV80 was the only MARV-infected bat with a detectable MARV-neutralizing titer (20) (**Figure 4J**).

At the 28 DPI necropsies, we cryopreserved spleens from EBOV-infected bats to evaluate lymphocyte populations compared to healthy controls with flow cytometry. The proportion of total T (CD3^+^) and B (CD79a^+^) lymphocytes in EBOV-infected bats (n=4) did not differ significantly from the healthy controls (n=8), although the proportion of T lymphocytes was elevated in spleens of bats infected with EBOV (**Figure 4K**). EBOV-infected bats had a distinct EBOV GP binding B cell population (mean 0.15%) compared to the healthy controls (**Figure 4L**).

Early B cell responses are characterized by the proliferation of plasmablasts and the formation of germinal centers in lymphoid tissues. Either process can lead to an expansion of specific B cell clonal lineages. Therefore, we analyzed the size of B cell clones in healthy controls (n=8) and EBOV-infected bats at 3-, 7-, and 28 DPI. Because MARV-infected bats did not mount a MARV GP-specific humoral response we did not evaluate their B cell clonal lineages. Among EBOV infected bats, the median count of B cell clones increased by 3 DPI and remained elevated until 7 DPI. The median count of B cell clones continued to increase from pre-infection levels to 28 DPI (**Supplementary Figure 4A**). However, due to significant individual-level variation at all time points, it was likely underpowered to detect statistical significance. This pattern was primarily observed among IgG sequences, and no clear pattern was observed among IgM sequences (**Supplementary Figure 4B**).

Next, we analyzed the frequency of VH and J gene segments. We restricted our analysis to only the most common VH gene family, the mouse homolog V5. Among IgG BCR sequences, the frequency of V5 sequences visually appeared to decrease at 3 DPI and return to pre-infection frequencies at 28 DPI (**Supplementary Figure 4C**). We analyzed the proportion of J segments focusing specifically on the most common J segment sequence, the mouse IGHJ3 gene homolog. Among IgM BCR sequences, the frequency of the J3 gene segment showed a statistically significant increase at 3 DPI (**Supplementary Figure 4D**). The J3 segment frequencies had returned to the level of healthy controls by 28 DPI.

### EBOV enters and replicates more efficiently than MARV in JFB cells

We previously published a study showing that JFB cells support the replication of both EBOV and MARV *in vitro*, and the growth of MARV seemed delayed compared to EBOV [43]. To determine if MARV growth is attenuated in JFB cells, we titrated the stocks of three EBOV strains (Kikwit, Makona C05, and Mayinga), and three MARV strains (Angola, Musoke, Ozolin), on Vero E6 and primary JFB uropatagium-derived fibroblasts (AjUFi_RML6). The titers of all the EBOV strains on JFB cells (Kikwit: 6.0 log_10_ TCID_50_/mL, Makona: 6.3 log_10_ TCID_50_/mL, Mayinga: 6.9 log_10_ TCID_50_/mL) were within 20-fold of the titers on Vero E6 cells (Kikwit: 6.6 log_10_ TCID_50_/mL, Makona: 7.6 log_10_ TCID_50_/mL, Mayinga: 7.5 log_10_ TCID_50_/mL), whereas the MARV titers were 2,000-32,622 fold lower on AjUFi_RML6 cells (Angola: 3.4 log_10_ TCID_50_/mL, Musoke: 2.2 log_10_ TCID_50_/mL, Ozolin: 3.6 log_10_ TCID_50_/mL) than on Vero E6 cells (Angola: 6.7 log_10_ TCID_50_/mL, Musoke: 6.7 log_10_ TCID_50_/mL, Ozolin: 7.1 log_10_ TCID_50_/mL) (**Figure 5A**).

**Figure 5.**
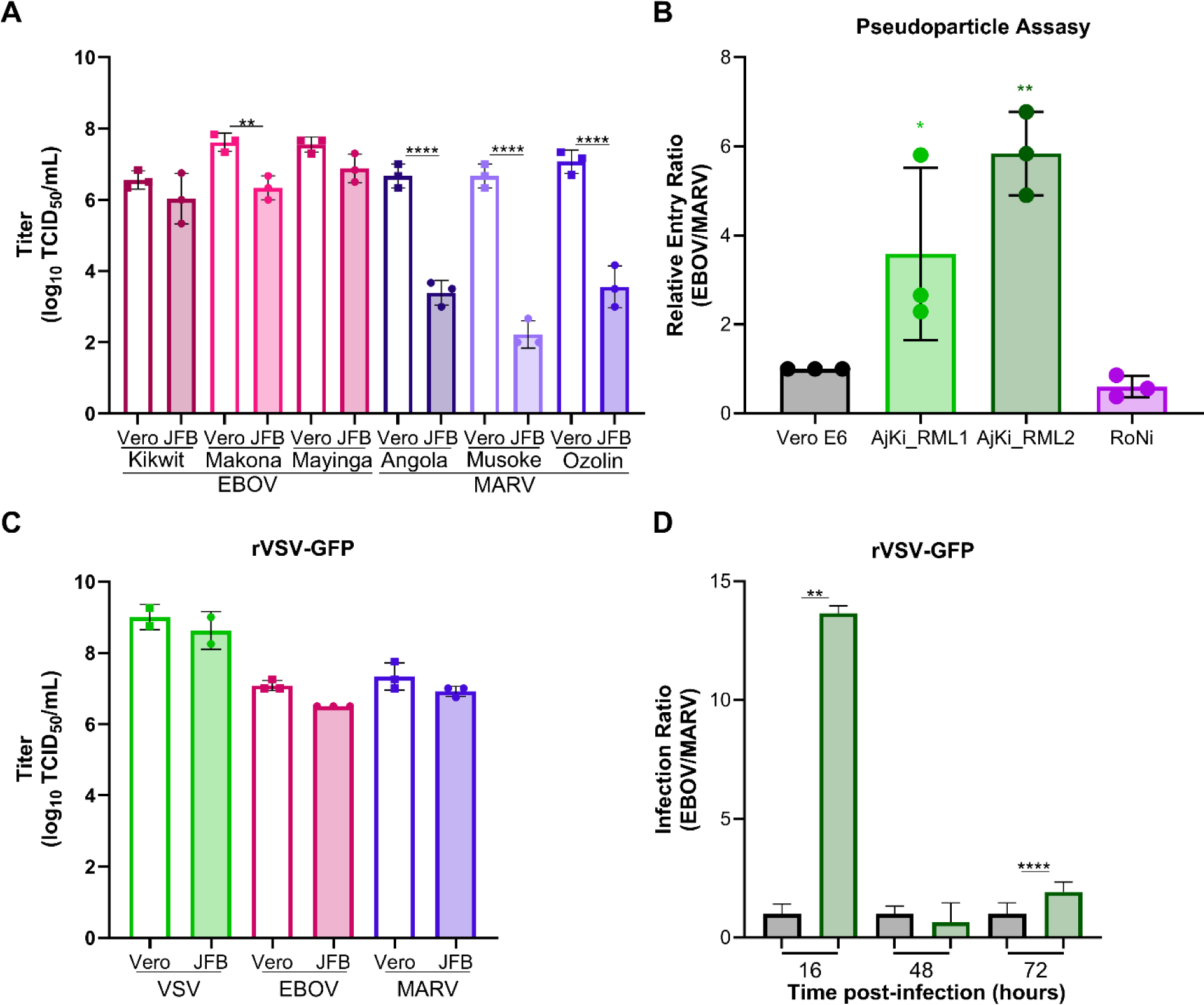
EBOV enters and replicates in JFB cells more efficiently than MARV. (A) Infectious titers of three EBOV strains (Kikwit, Makona C05, and Mayinga) and MARV strains (Angola, Musoke, and Ozolin) stocks measured on Vero E6 and JFB uropatagium-derived fibroblasts (AjUFi_RML6). Three independent stock vials were titrated on different days. The titer of each virus was compared on the two cell lines using a two-way ANOVA with Dunnett’s multiple test comparison. (B) Replication incompetent vesicular stomatitis virus (VSV) with glycoprotein gene replaced with green fluorescent protein pseudotyped with either EBOV or MARV glycoprotein (GP) were titrated on Vero E6, Jamaican fruit bat kidney (AjKi_RML1 and AjKi_RML2), or Egyptian rousette bat kidney (RoNi) cells. To determine the relative entry ratio, we normalized the ratio of EBOV/MARV infectious units for each cell line and divided by the ratio on Vero E6 cells. Three independent experiments were performed with four technical replicates each experiment. One-way ANOVA with Dunnett’s multiple test comparison. (C) Infectious titers of recombinant VSV expressing GFP (rVSVwt-GFP) or GFP plus EBOV-Mayinga or MARV-Ozolin GP were measured on Vero E6 and AjKi_RML2. Two (VSVwt) or three (EBOV and MARV GP) independent stock vials were titrated on two different passages of cells. The titer of each virus was compared on the two cell lines using a two-way ANOVA with Dunnett’s multiple test comparison. (D) Vero E6 and AjKi_RML2 cells were infected at multiplicity of infection (MOI) 0.005 for 1 hour and supernatants were collected at 16-, 48-, and 72 hours post-infection. The ratio of infectious titers for rVSV-, EBOV-GFP or rVSV-MARV-GFP was calculated for both cell lines and normalized to the ratio on Vero E6 cells (Replication kinetics shown in Supplementary Figure 5). Two independent experiments were performed with three technical replicates each experiment. The infection ratio at each time point was compared using a two-way ANOVA with Dunnett’s multiple test comparison.

To address this innate difference of viral growth on JFB cells, the contribution of viral entry was assessed. Non-replicating VSVΔGFP particles pseudotyped with either EBOV-Mayinga or MARV-Musoke GP, were titrated on Vero E6, AjKi_RML1, AjKi_RML2, and RoNi cells. The ratio of the VSVΔGFP-EBOV and VSVΔGFP-MARV titers were determined for each cell type and normalized to the Vero E6 ratio. For both JFB cell lines, VSVΔGFP-EBOV entered more efficiently than the VSVΔGFP-MARV (AjKi_RML1: 3.5-fold, AjKi_RML2: 5.8-fold) (**Figure 5B**). Although the VSVΔGFP-MARV trended toward more efficient entry on ERB cells, the difference was not significant (RoNi: 0.6-fold) (**Figure 5B**). To assess how the difference in entry may impact viral replication, we titrated VSVwild-type-GFP (VSVwt-GFP), VSV-EBOVmayinga-GFP (VSV-EBOV-GFP), or VSV-MARVozolin-GFP (VSV-MARV-GFP) on Vero E6 or AjKi_RML2 cells. Unlike WT EBOV and MARV, all three VSVs grew to comparable titers on both cell lines (**Figure 5C**). We also evaluated the replication kinetics of these VSVs on AjKi_RML2 cells (**Supplementary Figure 5A**) and Vero E6 cells (**Supplementary Figure 5B**). After normalization of the infection ratio of VSV-EBOV-GFP and VSV-MARV-GFP on AjKi_RML2 cells to the ratio on Vero E6 cells to account for intrinsic differences between the viruses’ replication, VSV-EBOV-GP grew to a titer 13.6-fold higher than VSV-MARV-GP at 16 HPI on JFB cells (**Figure 5D**). At 48- and 72 HPI the infection ratio of the two viruses was within 2-fold.

### EBOV antagonizes the JBF’s type I interferon pathway more efficiently than MARV

JFB and ERB cells were infected with EBOV-Mayinga or MARV-Ozolin to monitor replication kinetics and activation of the innate antiviral response. Consistent with previous results, MARV replication lagged from 24 (200-fold) through 72 (25-fold) HPI compared to EBOV in AjKi_RML2 cells, but both viruses reached similar titers at 96 (3-fold) and 120 (1.6-fold) HPI (**Figure 6A**). Although EBOV replicated to a higher titer compared to MARV at 72 HPI, MARV infection induced phosphorylation of STAT1 and STAT2 and expression of ISGs RIG-I and STAT1 as early as 72 HPI, whereas EBOV infection did not detectably activate the IFN-I signaling pathway through 120 HPI (**Figure 6B**). MARV infection induced expression of *Ifnb1* (**Figure 6C**) and ISG (**Figure 6D**) mRNAs 100- to 32,000-fold beginning at 72 HPI, and the same genes remained at mock levels in EBOV-infected cells. *Tnfa* expression was induced earlier in MARV-infected cells, but by 120 HPI levels were similar in both EBOV and MARV-infected cells (**Figure 6D**). Infection with either virus did not alter *Il6* expression more than 2-fold (**Figure 6E**). In ERB cells, EBOV and MARV replication kinetics were identical (**Supplementary Figure 6A**). MARV (48 HPI) infection induced *Ifnb* and ISG mRNAs earlier than EBOV (72 HPI) (**Supplementary Figures 6B, C**). RoNi cells infected with either virus increased expression of *Ifnb1* and ISG mRNAs to similar peak levels. Similar to the JFB cells, neither filovirus induced *Il6* but both induced *Tnfa* similarly in ERB cells (**Supplementary Figure 6D**). Despite similar peak levels of *Ifnb1* and ISG mRNAs, immunoblotting showed stronger activation of the IFN-I signaling pathway in MARV-infected cells (**Supplementary Figure 6E**).

**Figure 6.**
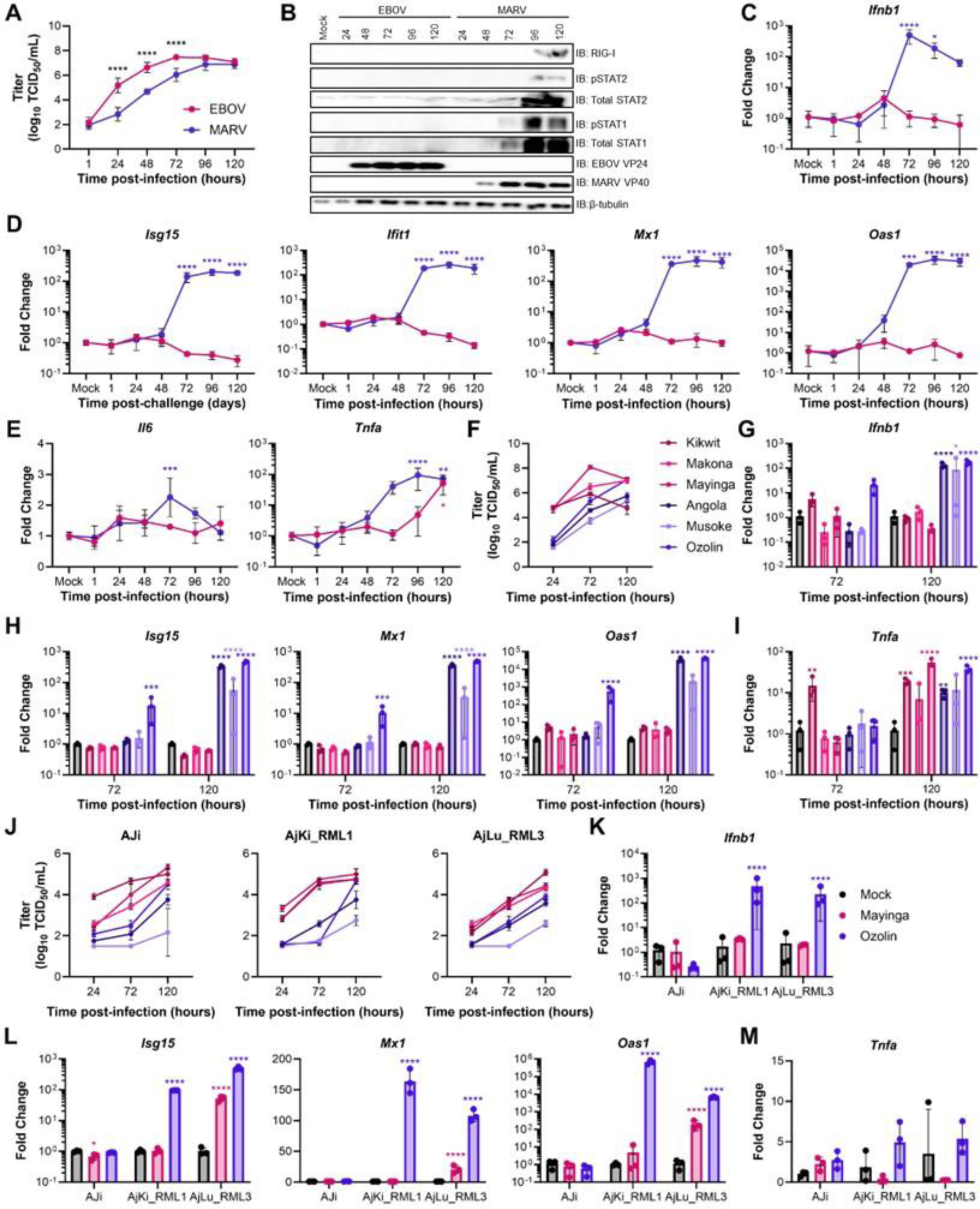
EBOV antagonizes JFB’s type I interferon pathway more efficiently than MARV. (A) Infectious titers of EBOV-Mayinga and MARV-Ozolin on Jamaican fruit bat kidney cells (AjKi_RML2) infected with multiplicity of infection (MOI) 0.1. Data presented as log_10_ transformed values. Data are from two independent experiments with three technical replicates each experiment. Two-way ANOVA with Sidak’s multiple test comparison to evaluate the difference between EBOV and MARV titer at each time point. (B) Immunoblot of AjKi_RML2 cells infected with EBOV or MARV at MOI 0.1. The presented panel is representative of three independent immunoblots. (C) RT-qPCR of *IFNb1* mRNA in AjKi_RML2 cells infected with EBOV or MARV at MOI 0.1. (D) RT-qPCR of interferon stimulated gene mRNAs in AjKi_RML2 cells infected with EBOV or MARV at MOI 0.1. (E) RT-qPCR of pro-inflammatory cytokine mRNAs in AjKi_RML2 cells infected with EBOV or MARV at MOI 0.1. (F) Infectious titers of three EBOV strains and three MARV strains on AjKi_RML2 cells infected with MOI 0.1. Data presented as log_10_ transformed values. Data are from one experiment conducted in triplicate. (G) RT-qPCR of *Infb1* mRNA in AjKi_RML2 cells infected with three different EBOV or MARV strains at MOI 0.1. (H) RT-qPCR of ISG mRNAs in AjKi_RML2 cells infected with three different EBOV or MARV strains at MOI 0.1. (I) RT-qPCR of pro-inflammatory cytokine *Tnf*a mRNA in AjKi_RML2 cells infected with three different EBOV or MARV strains at MOI 0.1. (J) Infectious titers of three EBOV strains and three MARV strains on Jamaican fruit bat immortalized kidney (AJi), primary kidney (AjKi_RML1), primary lung (AjLu_RML3) infected with MOI 0.1. Data presented as log_10_ transformed values. Data are from one experiment conducted in triplicate. (K) RT-qPCR of *Ifnb1* mRNA in Jamaican fruit bat cells infected with three EBOV Mayinga or MARV Ozolin strains at MOI 0.1. (L) RT-qPCR of ISG mRNAs in Jamaican fruit bat cells infected with three EBOV Mayinga or MARV Ozolin strains at MOI 0.1. (M) RT-qPCR of *Tnfa* mRNA in Jamaican fruit bat cells infected with three EBOV Mayinga or MARV Ozolin strains at MOI 0.1. (B-E; G-H; K-M) ΔC_T_ values were normalized to the average ΔC_T_ value of mock infected cells to calculate ΔΔC_T_. Fold change of each gene at each time point for EBOV and MARV-infected cells was compared to mock using a two-way ANOVA with Dunnett’s multiple test comparison.

To ensure that the intrinsic differences in replication and IFN-I signaling were not strain-specific, multiple JFB cell lines were infected with three different strains of EBOV and MARV. Importantly, back titration showed that the inoculation dose was within 5-fold for all the strains (**Supplementary Figure 6F**). All three EBOV strains replicated to higher titers than all three MARV strains at 24 and 72 HPI on AjKi_RML2 cells (**Figure 6F**). At 72 HPI only MARV-Ozolin induced expression *Ifnb1* and ISG mRNAs, but all three MARV strains induced these genes at 120 HPI (**Figures 6G, H**). None of the three EBOV strains induced expression of the evaluated IFN-I pathway genes. EBOV-Kikwit induced *Tnfa* at 72 HPI, and infection increased *Tnfa* 3-32 fold at 120 HPI regardless of strain (**Figure 6I**). Although the magnitude of the difference varied across cell lines, all EBOV strains replicated to higher titer than MARV strains on three additional JFB cell lines (Aji, AjKi_RML2, and AjLu_RML3) (**Figure 6J**). Because we observed similar patterns in host gene expression across the different virus strains on AjKi_RML2 cells and similar replication kinetics across all JFB cell lines, changes to host mRNAs were measured in the other JFB cell lines 120 HPI with EBOV-Mayinga and MARV-Ozolin. *Ifnb1* was induced in AjKi_RML1 and AjLu_RML3 cells infected with MARV but not EBOV (**Figure 6K**). In AjKi_RML1 cells expression of all ISGs increased in MARV-infected cells compared with mock while EBOV infection did not alter ISG mRNA levels (**Figure 6L**). ISG mRNAs increased in both EBOV and MARV infected AjLu_RML3 cells, but the magnitude of expression was higher in MARV-infected cells than EBOV-infected cells despite replication to lower titer throughout infection (**Figures 6J, L**). AJi cells express very low levels of endogenous STAT1 compared to the other cell lines, which causes AJi cells to have a higher threshold for activation of the IFN-I signaling pathway. In AJi cells, levels of *Ifnb1* and ISG mRNAs did not change in either EBOV- or MARV-infected cells relative to mock (**Figure 6K, L**). TNF mRNA levels did not differ in any of the cell lines infected with either virus (**Figure 6M**).

Replication kinetics with all the EBOV and MARV strains were conducted on human and ERB cell lines to confirm that the differences were not virus intrinsic. Overall, the titers of the EBOV and MARV strains were similar throughout the time course for human (Huh7) (**Supplementary Figure 6G**) and two ERB kidney (RoNi and RASKM) cell lines (**Supplementary Figure 6H**). In contrast to what is observed on JFB cells, all three MARV strains grew to a higher titer on ERB lung cells (RaLu) than all three EBOV strains at 120 HPI (**Supplementary Figure 6H**). *Ifnb1* (**Supplementary Figure 6I**), ISGs (**Supplementary Figure 6J**), and *Tnfa* (**Supplementary Figure 6K**) mRNAs were elevated in RoNi cells infected with each of the viruses at 120 HPI.

### Inability of MARV to antagonize IFN-I signaling partially contributes to attenuated replication in JFB cells

Due to the activation of the IFN-I pathway in MARV-, but not EBOV-infected cells, the capacity of the viruses to antagonize IFN-I signaling was compared. MARV VP40 and EBOV VP24 block the activation of the IFN-I response pathway at different points. MARV VP40 prevents the JAK1 phosphorylation and downstream phosphorylation of STAT1 [44, 45], whereas EBOV VP24 interacts with the NPI-1 subfamily of karyopherin alpha proteins to preclude pSTAT1’s nuclear translocation [46–50] (**Figure 7A**). To measure the functionality of EBOV and MARV IFN-I signaling antagonists, Huh7 cells were infected with EBOV-Mayinga or MARV-Ozolin at MOI 1.5 for 24 hours. Cells were stimulated with 100 ng/mL of recombinant Chiroptera IFN-β for 30 minutes prior to cytoplasmic-nuclear fractionation. As expected, pSTAT1 was present in both fractions of mock infected cells treated with IFN-β (**Supplementary Figure 7A**). Cytoplasmic pSTAT1 levels were similar in mock and EBOV-infected cells treated with IFNβ, but nuclear pSTAT1 was 50% lower in EBOV-infected cells (**Supplementary Figures 7A, B**). Both cytoplasmic and nuclear pSTAT1 was reduced over 90% in MARV-infected cells compared to mock (**Supplementary Figures 7A, B**). Huh7 cells cannot produce IFN-β, thus the experiment does not require an IFN-I induction blockade after infection. Because the experiment requires a high amounts of virus replication to ensure expression of viral protein and IFN-I cannot be induced prior to the addition of exogenous IFN-β, we confirmed that the TBK1 and IKKε specific inhibitor BAY-985 [51] blocked IFN-I induction in AjKi_RML2 cells after transfection with 500 ng/mL of high molecular weight poly(I:C) (**Supplementary Figures 8A, B**). AjKi_RML2 cells were infected with EBOV-Mayinga or MARV-Ozolin at MOI 1.5 and treated with 25 nM BAY-985 for 24 hours prior to the addition of 100 ng/mL Chiroptera IFNβ for 30 minutes. After IFNβ treatment, EBOV-infected cells had elevated cytoplasmic pSTAT1 and significantly reduced nuclear pSTAT1 (90% inhibition) compared to mock infected cells (**Figures 7B, C**). Although the levels of cytoplasmic pSTAT1 was decreased significantly (25% inhibition), the amount of nuclear pSTAT1 did not differ between MARV- and mock-infected cells after IFN-β addition (**Figures 7B, C**).

**Figure 7.**
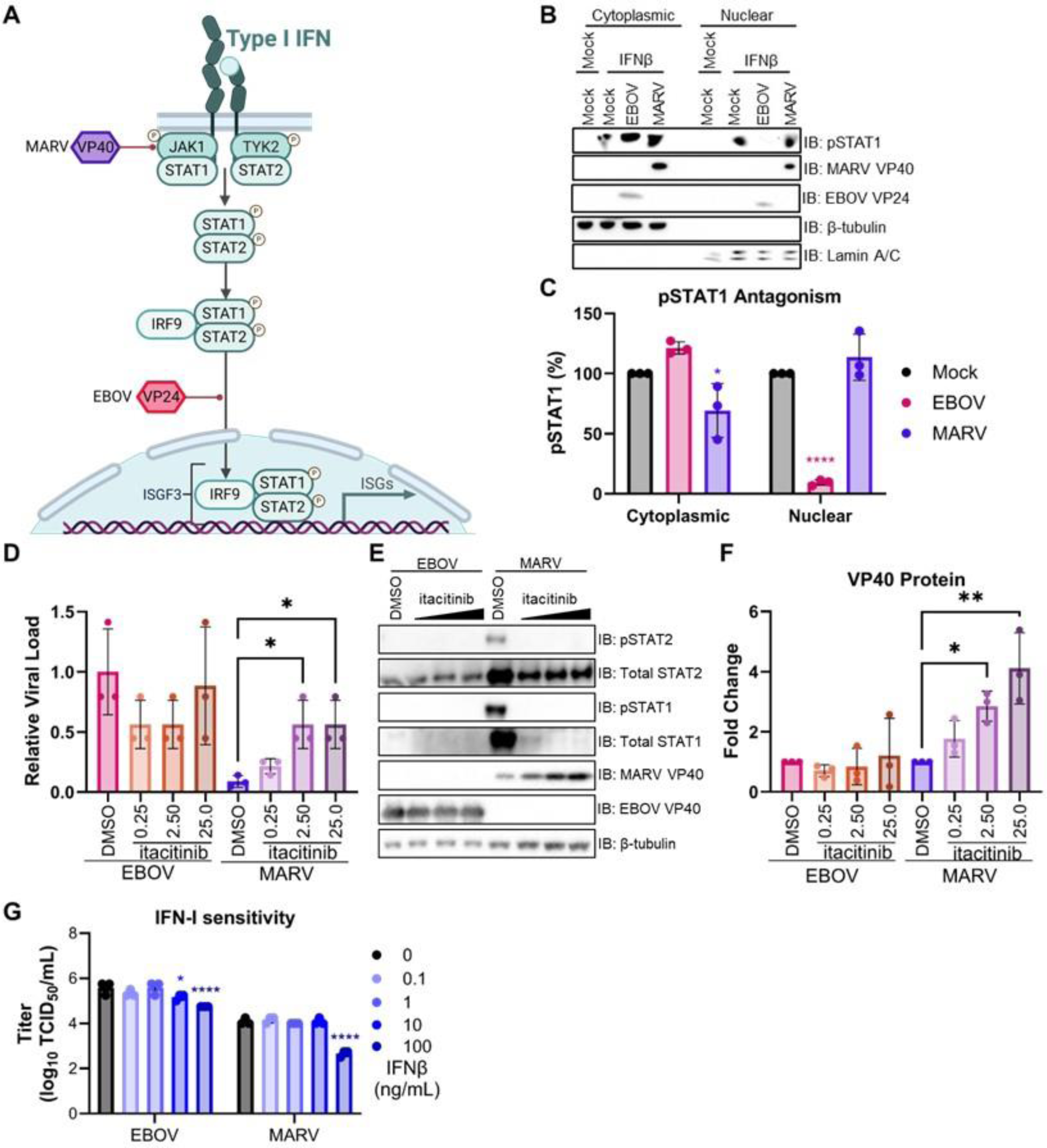
Inability of MARV to antagonize IFN-I signaling partially contributes to attenuated replication. (A) Schematic mechanisms of EBOV VP24 and MARV VP40 antagonism of the type-I interferon signaling pathway. Image created using Biorender. (B) Immunoblot of cytoplasmic and nuclear fractions of Jamaican fruit bat kidney cells, AjKi_RML2, infected with MOI 1.5 of EBOV-Mayinga or MARV-Ozolin for 24 hours prior to the addition of exogenous recombinant Chiroptera IFN-β (100ng/mL) for 30 minutes. Cells were treated with type I interferon induction inhibitor BAY-985 (25 nM) at the time of infection. The presented panel is representative of three independent immunoblots. (C) Quantification of phosphorylated STAT1 in the cytoplasmic and nuclear fractions. Normalized expression was determined using the following formula: (AUC pSTAT1)/(AUC β-tubulin (cytoplasmic) or Lamin A/C (nuclear)). To calculate relative expression, the normalized sample value is divided by the normalized expression in uninfected cells treated with IFNβ. Three independent experiments were performed. Percentage of pSTAT1 in each of the fractions for EBOV and MARV-infected cells was compared mock using a two-way ANOVA with Dunnett’s multiple test comparison. (D) AjKi_RML2 cells were infected with MOI 0.1 EBOV-Mayinga or MARV-Ozolin for 1 hour and treated with vehicle control DMSO or itacitinib (0.25-25 nM) immediately after infection. Fresh media with inhibitor was replaced at 24- and 48-hours post infection. Supernatants collected at 72 hours post-infection were titrated. To calculate the relative viral load, the viral titers were normalized to the average titer of EBOV-infected cells treated with DMSO. The experiment was performed in triplicate. The effect of itacitinb on EBOV and MARV replication was determined using a one-way ANOVA with Dunnett’s multiple test comparison. (E) Immunoblot showing the effect of itacitinib treatment on the activation of the immune response and VP40 expression. The panel is representative of two independent immunoblots. (F) Quantification of VP40 for the immunoblot presented in panel D. Normalized expression was determined using the following formula: (AUC VP40)/(AUC β-tubulin). To calculate relative expression, the normalized sample value is divided by the normalized expression in DMSO-treated cells infected with either EBOV or MARV. (G) Infectious titers of EBOV-Mayinga or MARV-Ozolin at 48 hours post-infection. AjKi_RML2 cells were treated with recombinant Chiroptera IFN-β in 10-fold serial dilution (0-100 ng/mL) for 24 hours prior to infection with MOI 0.1. Data presented as log_10_ transformed values. The experiment was performed in triplicate. To test a change in viral titer at each dose compared to mock treated cells, we performed a two-way ANOVA with Dunnett’s multiple test comparison.

To evaluate the importance of MARV VP40 impaired signaling to viral replication, IFN-I signaling was inhibited. The effectiveness of JAK1 inhibitor itacitinib was confirmed first in AjKi_RML2 cells stimulated with 100 ng/mL IFNβ for 30 minutes (**Supplementary Figure 8B**) or transfected with poly(I:C) for 16 hours (**Supplementary Figure 8C**). AjKi_RML2 cells were infected with EBOV-Mayinga or MARV-Ozolin at a MOI 0.1 for 1 hour before addition of itacitinib (0.25-25 nM) in fresh medium. Medium with inhibitor was replaced fully at 24 and 48 HPI, and viral titer in the supernatant at 72 HPI was measured. The MARV titer increased in a dose-dependent manner, but itacitinib treatment did not impact EBOV replication relative to infected cells treated with the vehicle control, DMSO (**Figure 7D**). In correlation with the increased MARV titer, elevated VP40 protein expression was itacitinib dose-dependent (**Figures 7E, 7F**). EBOV VP40 protein levels remained relatively constant and were independent of the itacitinib concentration (**Figures 7E, 7F**).

The IFN-I sensitivity of EBOV and MARV in JFB and ERB cells was tested to assess whether bat ISGs differ in their capacity to block filovirus replication. AjKi_RML2 and RoNi cells were treated with IFN-β (0-100 ng/mL) for 24 hours prior to infection with MOI 0.1 of EBOV-Mayinga or MARV-Ozolin. At 48 HPI, the IFN-I sensitivity of EBOV and MARV was similar on JFB cells (**Figure 7G**), but EBOV is more sensitive to IFN-I than MARV in ERB cells (**Supplementary Figure 8D**). IFN-β treatment resulted in the dose-dependent increase of ISG mRNAs in JFB (**Supplementary Figure 8E**) and ERB cells (**Supplementary Figure 8F**) as expected.

## Discussion

Despite the mounting evidence that bats naturally host orthoebolaviruses, the lack of a full-length genome sequence or isolate for all species that cause disease in humans limits our understanding of which bat species may serve as reservoir hosts. In support of the hypothesis that bat species naturally host EBOV, we demonstrated that JFB are robustly susceptible to EBOV. Like the recently reported Angolan free-tailed bat model [52], we observed EBOV replication and oral shedding while MARV infection is transient and quenched rapidly. In contrast, ERBs are susceptible to MARV but not orthoebolaviruses [2]. These results suggest that bat species have inherent differences in their susceptibility to different filovirus species. Future studies evaluating JFB infections with other filovirus species will enhance understanding of the within-host factors that dictate virus-host complementarity.

The correlation of the *in vitro* data with the observed JFB infection outcomes with EBOV and MARV supports that phylogenomics may be useful in predicting which bat species may support a given filovirus species. In support of the decreased entry of MARV pseudoparticles and VSV-MARV-GFP compared to the EBOV GP particles, JFB NPC1, the receptor for EBOV and MARV [53–55], has amino acid differences in the loops that interact with filovirus GP. Future studies using recombinant, mutant JFB NPC1 or filovirus GPs can be used to evaluate the contribution of specific residues in entry. Compared to EBOV, MARV inefficiently blocked the IFN-I pathway in JFB cells. Although the phenotype should be investigated in dendritic cells and macrophages at timepoints following early infection, the impaired immune antagonism of MARV likely contributes to the poor replication and dissemination *in vivo*. Since MARV replication significantly lags compared to EBOV (2 log_10_ TCID_50_/mL) *in vitro* before the IFN-I response is induced, budding, replication, and/or transcription may be attenuated. Evaluation of replication and transcription in JFB cells using the minigenome system is not technically feasible currently, but we are actively working to improve transfection protocols to allow direct evaluation of polymerase functional differences between EBOV and MARV. Adaptation of MARV to JFBs could reveal key mutations needed to enable susceptibility. Importantly, ERBs, despite supporting MARV but not orthoebolaviruses *in vivo*, do not have differential replication kinetics *in vitro*. However, we observed that MARV is insensitive to IFN-I in ERB cells relative to EBOV. This suggests that MARV-ERB co-evolution may have enabled MARV to evade the antagonistic functions of ISGs. Different stages in the filovirus life cycle are likely critical in determining virus-host complementarity among bat species. Studying the kinetics of filoviruses in cell lines, preferably representative of different bat families, can contribute knowledge of key innate host factors that influence host susceptibility.

Like other bat-filovirus infection models [1, 2, 52], EBOV-infected JFBs did not experience morbidity or mortality. In the acute phase, we observed minor-to-mild signs of disease including hypothermia and weight loss (≤5%) in some of the EBOV-infected bats. The infection did not result in any gross lesions and only minor histopathologic changes at the time points evaluated. Unlike infections in dead-end hosts, which can present with hemorrhagic disease [41, 42], EBOV infection did not result in lymphopenia or neutrophilia in JFBs. EBOV infection increased the number of circulating monocytes suggesting induction of an early inflammatory response. Akin to other bat-filovirus models [5, 56], JFBs mounted a potent IFN-I response while changes in expression of pro-inflammatory cytokine genes was mild. Although the underlying mechanisms in the JFB-EBOV system still need to be evaluated, the lack of a robust inflammatory cytokine response could be tied to differences in inflammasome function [57–60] or STING activation [61] observed in other bat species. Guito and co-authors demonstrated in the ERB-MARV model that dexamethasone-induced immune suppression bolstered MARV replication and shedding and was associated with gross lesions [62]. Although the EBOV-infected JFBs all survived infection, we hypothesize that inoculation with a higher dose or immune suppressing the animal prior to infection could result in a lethal outcome for some individuals.

The JFB-EBOV model is amenable to investigating the factors that influence viral shedding. Infectious EBOV in oral swabs peaked at 6 DPI. These data resemble those reported for the MARV-ERB model in both magnitude and timing [1]. The variation in the amount of virus shed was substantial, suggesting within-host factors could influence shedding [63, 64]. Manipulating the bats’ diet to mimic nutritional stress or altering climatic conditions could be used to elucidate the impact of environmental stressors on shedding kinetics as well as ISG and cytokine expression. Pregnancy infection models could also be important to address the influence of reproductive status on infection outcome and vertical transmission.

Establishment of this JFB-EBOV model allows us to interrogate a plethora of open questions in the field. A primary area of concern is the contribution of the host’s immune system in controlling infection. Early in infection, dendritic cells and/or macrophages provide an environment for filoviruses to replicate before dissemination [41, 42, 65]. Future studies will investigate the early innate responses at the inoculated site in response to EBOV and MARV infection to evaluate host factors that may predict infection outcome. Compared to mice that tolerate infection with mouse-adapted EBOV [66], infected JFBs have a strong antiviral transcriptional profile in the liver 3 DPI. Beyond the innate response, the adaptive immune system is critical for controlling infection. In dead-end hosts, filoviruses can infect T lymphocytes and cause cell death through abortive infection [67]. Whether or not filoviruses infect bat T cells remains unknown and could offer insight into differences between lethal and non-lethal infection. Comparing the infections between EBOV and MARV can also provide information of the establishment of an adaptive, humoral response in bats as we observed a MARV GP-specific antibody response in only one of the four infected bats. In the Angolan free-tail bat-EBOV model, vertical transmission was documented [52]. Although we detected EBOV RNA in ovary and testis samples collected at 3 and 7 DPI, whether vertical transmission can be achieved in the EBOV-JFB model needs to be investigated.

Another prevalent question in the field of bat-associated viruses is whether virus persists and recrudesces when a chronically infected bat encounters different stressors. At 28 DPI, we detected EBOV RNA in the spleen at a similar magnitude to 3 and 7 DPI. Due to insufficient sample, we were unable to titrate spleen homogenates collected at 28 DPI and cannot determine whether the virus present was infectious or not. Future studies can address the potential for persistence of infectious virus. In a transmission study using the ERB-MARV model, Schuh and co-authors observed transmission after the acute phase of infection which suggests viral persistence may be a feature of filoviruses [3]. Further, human survivors of Ebola virus disease (EVD) can have persistent viral RNA and shed infectious virus [68–71].

Overall, JFBs are a valuable model for interrogating filovirus biology. Differential susceptibility to EBOV and MARV offers an opportunity to explore the intricacies of virus-host relationships. Support of EBOV replication with disseminated infection and shedding of infectious virus enables the evaluation of both biotic and abiotic factors that regulate shedding and transmission.

## Acknowledgements

We would like to thank the animal caretakers for their assistance with animal husbandry. This study was supported by the Intramural Research Program of NIAID, NIH.

## Author Contributions

SvT, JRP designed the studies.

SvT, JRP, RJF, SG, TB, AG, JES, AC, LM, DEC, CAF, JCR, AW, CC, JL, CS, GS, JP-S, performed the experiments.

SvT, JRP, JCR, LM, DC, CC, JL, CM, SA analyzed results.

YH, TS, LK, and AM provided essential reagents and support.

SvT, VM wrote the manuscript.

All co-authors reviewed the manuscript and approved this version.

---

**Supplementary Table 1.**
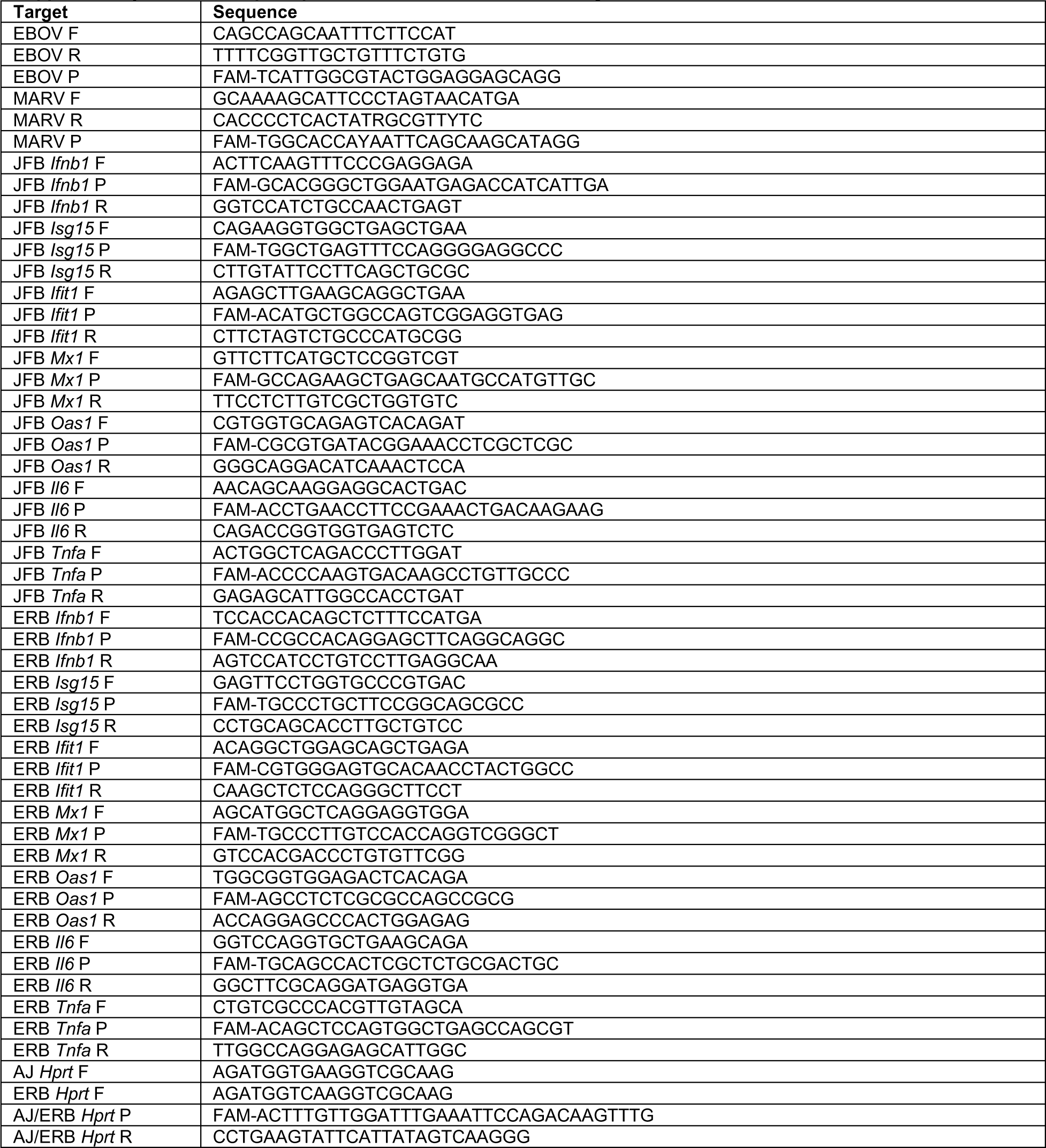
RT-qPCR primers for JFB and ERB immune genes and viruses.

**Supplementary Table 2.** Hematology of EBOV and MARV infected JFBs.

| Study Day | Animal ID | RBC (M/uL) | WBC (K/uL) | HGB (g/dL) | NEUT# (K/uL) | NEUT% (%) | HCT (%) | LYMPH# (K/uL) | LYMPH% (%) | MCV (fL) | MONO# (K/uL) | MONO% (%) | MCH (pg) | EOS# (K/uL) | EOS% (%) | MCHC (g/dL) | BASO# (K/uL) | BASO% (%) | RDW SD (fL) | RDW CV (%) | PLT (K/uL) | MPV (fL) | RET# (K/uL) | RET% (%) |
| --- | --- | --- | --- | --- | --- | --- | --- | --- | --- | --- | --- | --- | --- | --- | --- | --- | --- | --- | --- | --- | --- | --- | --- | --- |
| -7 | EBO01 | 14.36 | 10.94 | 19.6 | 5.02 | 45.9 | 48.9 | 5.15 | 47.1 | 34.1 | 0 | 0 | 13.6 | 0 | 0 | 40.1 | 0.77 | 7 | 31 | 37.1 | 456 | 9.3 | 110.6 | 0.77 |
| 3 | EBO01 | 13.63 | 5.19 | 18.9 | 2.68 | 51.6 | 46.6 | 2.22 | 42.8 | 34.2 | 0.05 | 1 | 13.9 | 0 | 0 | 40.6 | 0.24 | 4.6 | 29.3 | 35.6 | 752 | 12 | 205.8 | 1.51 |
| -8 | EBO02 | 12.96 | 5.73 | 17.7 | 3.06 | 53.4 | 42.5 | 2.67 | 46.6 | 32.8 | 0 | 0 | 13.7 | 0 | 0 | 41.6 | 0 | 0 | 27.1 | 33.8 | 420 | 10.6 | 134.8 | 1.04 |
| 3 | EBO02 | 12.23 | 5.29 | 16.7 | 1.86 | 35.2 | 41.8 | 2.88 | 54.4 | 34.2 | 0.04 | 0.8 | 13.7 | 0 | 0 | 40 | 0.51 | 9.6 | 26.2 | 31.3 | 448 | 13 | 332.7 | 2.72 |
| -8 | EBO07 | 12.93 | 10.45 | 17.5 | 1.78 | 17 | 43 | 8.42 | 80.6 | 33.3 | 0.01 | 0.1 | 13.5 | 0 | 0 | 40.7 | 0.24 | 2.3 | 26.6 | 33.5 | 425 | 9.7 | 250.8 | 1.94 |
| 3 | EBO07 | 13.25 | 7.16 | 17.5 | 0.49 | 6.8 | 44.5 | 6.52 | 91.1 | 33.6 | 0.06 | 0.8 | 13.2 | 0 | 0 | 39.3 | 0.09 | 1.3 | 26.1 | 32.8 | 613 | 12 | 382.9 | 2.89 |
| -8 | EBO08 | 13.89 | 10.99 | 18.7 | 8.5 | 77.3 | 43.4 | 2.22 | 20.2 | 31.2 | 0.01 | 0.1 | 13.5 | 0 | 0 | 43.1 | 0.26 | 2.4 | 26.2 | 37.9 | 521 | 8.8 | 123.6 | 0.89 |
| 3 | EBO08 | 12.34 | 7.6 | 16.4 | 5.35 | 70.4 | 39.5 | 2.07 | 27.2 | 32 | 0.09 | 1.2 | 13.3 | 0.01 | 0.1 | 41.5 | 0.08 | 1.1 | 24.9 | 33.4 | 451 | 12.9 | 246.8 | 2 |
| -8 | EBO03 | 14.52 | 3.03 | 20.2 | 2.34 | 77.2 | 49.5 | 0.69 | 22.8 | 34.1 | 0 | 0 | 13.9 | 0 | 0 | 40.8 | 0 | 0 | 29.4 | 36.8 | 547 | 10.8 | 88.6 | 0.61 |
| 7 | EBO03 | 12.07 | 10.21 | 16.5 | 7.02 | 68.7 | 41.2 | 3.14 | 30.8 | 34.1 | 0.05 | 0.5 | 13.7 | 0 | 0 | 40 | 0 | 0 | 25.6 | 30.7 | 799 | 12.1 | 144.8 | 1.2 |
| -7 | EBO04 | 13.42 | 10.56 | 19.2 | 2.56 | 24.2 | 49.1 | 8 | 75.8 | 36.6 | 0 | 0 | 14.3 | 0 | 0 | 39.1 | 0 | 0 | 26.7 | 30.7 | 537 | 10.4 | 95.3 | 0.71 |
| 7 | EBO04 | 12.78 | 14.18 | 18.3 | 8.42 | 59.4 | 47.5 | 5.56 | 39.2 | 37.2 | 0.09 | 0.6 | 14.3 | 0 | 0 | 38.5 | 0.11 | 0.8 | 26.5 | 30 | 733 | 11.1 | 337.4 | 2.64 |
| -8 | EBO09 | 13.93 | 5.18 | 18.6 | 2.36 | 45.6 | 46.6 | 2.82 | 54.4 | 33.5 | 0 | 0 | 13.4 | 0 | 0 | 39.9 | 0 | 0 | 26.8 | 34.3 | 889 | 9.5 | 79.4 | 0.57 |
| 7 | EBO09 | 12.45 | 10.83 | 16.4 | 4.77 | 44 | 41 | 5.8 | 53.6 | 32.9 | 0.02 | 0.2 | 13.2 | 0 | 0 | 40 | 0.24 | 2.2 | 24.8 | 31.3 | 902 | 11.1 | 146.9 | 1.18 |
| -8 | EBO10 | 14.12 | 4.85 | 18.7 | 2.69 | 55.5 | 47.1 | 2.16 | 44.5 | 33.4 | 0 | 0 | 13.2 | 0 | 0 | 39.7 | 0 | 0 | 28.3 | 35.8 | 540 | 9.1 | 89 | 0.63 |
| 7 | EBO10 | 12.91 | 12.72 | 16.9 | 3.89 | 30.6 | 42.6 | 8.75 | 68.8 | 33 | 0.08 | 0.6 | 13.1 | 0 | 0 | 39.7 | 0 | 0 | 27.1 | 33.3 | 772 | 11.8 | 227.2 | 1.76 |
| -8 | EBO05 | 13.76 | 4.63 | 18.9 | 2.14 | 46.2 | 45.1 | 2.37 | 51.2 | 32.8 | 0 | 0 | 13.7 | 0 | 0 | 41.9 | 0.12 | 2.6 | 27.8 | 36.4 | 33 | 6.7 | 60.5 | 0.44 |
| 28 | EBO05 | 12.91 | 6.97 | 17.2 | 2.87 | 41.2 | 42.3 | 4.09 | 58.7 | 32.8 | 0.01 | 0.1 | 13.3 | 0 | 0 | 40.7 | 0 | 0 | 26.5 | 33.5 | 943 | 9.9 | 370.5 | 2.87 |
| -8 | EBO06 | 14.8 | 5.78 | 20.1 | 2.86 | 49.5 | 47.6 | 2.6 | 45 | 32.2 | 0 | 0 | 13.6 | 0 | 0 | 42.2 | 0.32 | 5.5 | 28.9 | 39.4 | 511 | 8.9 | 97.7 | 0.66 |
| 28 | EBO06 | 13.47 | 5.63 | 17.8 | 1.5 | 26.6 | 44.4 | 4.04 | 71.8 | 33 | 0.02 | 0.4 | 13.2 | 0 | 0 | 40.1 | 0.07 | 1.2 | 27.7 | 34.9 | 950 | 10.6 | 281.5 | 2.09 |
| -8 | EBO11 | 11.74 | 11.51 | 15.9 | 9.42 | 81.8 | 40.8 | 2.08 | 18.1 | 34.8 | 0.01 | 0.1 | 13.5 | 0 | 0 | 39 | 0 | 0 | 25 | 29.3 | 816 | 10.5 | 179.6 | 1.53 |
| 28 | EBO11 | 11.84 | 12.45 | 16.4 | 4.63 | 37.1 | 42.1 | 7.8 | 62.7 | 35.6 | 0.02 | 0.2 | 13.9 | 0 | 0 | 39 | 0 | 0 | 26 | 29 | 1163 | 11.3 | 440.4 | 3.72 |
| -8 | EBO12 | 12.57 | 8.31 | 18.4 | 1.66 | 20 | 47.4 | 6.65 | 80 | 37.7 | 0 | 0 | 14.6 | 0 | 0 | 38.8 | 0 | 0 | 25.7 | 28.4 | 790 | 10.1 | 206.1 | 1.64 |
| 28 | EBO12 | 11.97 | 10.03 | 17.3 | 2.01 | 20 | 43.9 | 7.75 | 77.3 | 36.7 | 0.01 | 0.1 | 14.5 | 0 | 0 | 39.4 | 0.26 | 2.6 | 25.1 | 28.1 | 621 | 12.2 | 397.4 | 3.32 |
| -7 | MARV01 | 14.05 | 3.32 | 19.3 | 1.93 | 58.1 | 47.3 | 1.39 | 41.9 | 33.7 | 0 | 0 | 13.7 | 0 | 0 | 40.8 | 0 | 0 | 27.1 | 34.9 | 521 | 10.2 | 61.8 | 0.44 |
| 3 | MARV01 | 11.98 | 3.02 | 16.4 | 1.33 | 44 | 41.6 | 1.67 | 55.3 | 34.7 | 0.02 | 0.7 | 13.7 | 0 | 0 | 39.4 | 0 | 0 | 24.3 | 28.6 | 684 | 12 | 135.4 | 1.13 |
| -7 | MARV02 | 2.49 | ---- | 3.3 | ---- | ---- | 6.2 | ---- | ---- | 24.9 | ---- | ---- | 13.3 | ---- | ---- | 53.2 | ---- | ---- | 20.9 | 22.4 | 81 | 9.8 | 15.2 | 0.61 |
| 3 | MARV02 | 13.47 | 5.36 | 18 | 1.96 | 36.5 | 44.4 | 3.22 | 60.1 | 33 | 0.02 | 0.4 | 13.4 | 0 | 0 | 40.5 | 0.16 | 3 | 27.7 | 34.7 | 725 | 11.4 | 255.9 | 1.9 |
| -7 | MARV07 | 12.54 | 24 | 17.4 | 5.64 | 23.4 | 39.7 | 16.72 | 69.7 | 31.7 | 0.02 | 0.1 | 13.9 | 0 | 0 | 43.8 | 1.62 | 6.8 | 24.1 | 34.7 | 333 | 8.8 | 267.1 | 2.13 |
| 3 | MARV07 | 13.61 | 20.56 | 18.3 | 4.27 | 20.9 | 44.1 | 16.06 | 78.1 | 32.4 | 0.17 | 0.8 | 13.4 | 0.01 | 0 | 41.5 | 0.05 | 0.2 | 24.8 | 34.3 | 221 | 10.3 | 371.6 | 2.73 |
| -7 | MARV08 | 13.26 | 10.23 | 17.5 | 5.41 | 52.9 | 42.6 | 4.82 | 47.1 | 32.1 | 0 | 0 | 13.2 | 0 | 0 | 41.1 | 0 | 0 | 26.3 | 35.4 | 496 | 9.6 | 110.1 | 0.83 |
| 3 | MARV08 | 11.82 | 3.37 | 15.7 | 1.88 | 55.8 | 39.4 | 1.43 | 42.4 | 33.3 | 0.01 | 0.3 | 13.3 | 0 | 0 | 39.8 | 0.05 | 1.5 | 27.2 | 31.6 | 604 | 10.7 | 450.3 | 3.81 |
| -7 | MARV03 | 12.98 | 7.45 | 18 | 3.93 | 52.8 | 44.1 | 3.52 | 47.2 | 34 | 0 | 0 | 13.9 | 0 | 0 | 40.8 | 0 | 0 | 26.4 | 32.5 | 631 | 10.3 | 106.4 | 0.82 |
| 7 | MARV03 | 11.77 | 6.98 | 16.1 | 3.05 | 43.6 | 40.6 | 3.85 | 55.2 | 34.5 | 0.04 | 0.6 | 13.7 | 0 | 0 | 39.7 | 0.04 | 0.6 | 24.5 | 29.2 | 834 | 12.2 | 258.9 | 2.2 |
| -7 | MARV04 | 12.94 | 6.49 | 18.7 | 3.28 | 50.6 | 44.5 | 2.85 | 43.9 | 34.4 | 0 | 0 | 14.5 | 0 | 0 | 42 | 0.36 | 5.5 | 26 | 32.1 | 635 | 9.5 | 116.5 | 0.9 |
| 7 | MARV04 | 12.24 | 3.87 | 17.3 | 1.44 | 37.2 | 44.1 | 2.43 | 62.8 | 36 | 0 | 0 | 14.1 | 0 | 0 | 39.2 | 0 | 0 | 26.1 | 28.9 | 991 | 10.7 | 308.4 | 2.52 |
| -7 | MARV09 | 13.83 | 14.17 | 19.4 | 4.67 | 32.9 | 47.7 | 8.68 | 61.3 | 34.5 | 0.01 | 0.1 | 14 | 0 | 0 | 40.7 | 0.81 | 5.7 | 28.9 | 34.6 | 812 | 10 | 113.4 | 0.82 |
| 7 | MARV09 | 12.93 | 15.42 | 18.2 | 9.85 | 63.9 | 45 | 5.51 | 35.7 | 34.8 | 0.06 | 0.4 | 14.1 | 0 | 0 | 40.4 | 0 | 0 | 27.9 | 32.1 | 1065 | 11.5 | 350.4 | 2.71 |
| -7 | MARV10 | 14.86 | 5.91 | 19.3 | 2.52 | 42.7 | 45.6 | 3.14 | 53.1 | 30.7 | 0 | 0 | 13 | 0 | 0 | 42.3 | 0.25 | 4.2 | 28.2 | 40.4 | 574 | 9.5 | 173.9 | 1.17 |
| 7 | MARV10 | 14.49 | 9.65 | 18.7 | 7.92 | 82.1 | 45.2 | 1.71 | 17.7 | 31.2 | 0.02 | 0.2 | 12.9 | 0 | 0 | 41.4 | 0 | 0 | 28.2 | 38.9 | 857 | 10.2 | 336.2 | 2.32 |
| -7 | MARV05 | 12.03 | 3.01 | 17.3 | 1.63 | 54.2 | 43.7 | 1.18 | 39.2 | 36.3 | 0 | 0 | 14.4 | 0 | 0 | 39.6 | 0.2 | 6.6 | 26.3 | 28.5 | 446 | 8.9 | 156.4 | 1.3 |
| 28 | MARV05 | 11.8 | 3.78 | 17.6 | 0.96 | 25.4 | 46.2 | 2.8 | 74.1 | 39.2 | 0.02 | 0.5 | 14.9 | 0 | 0 | 38.1 | 0 | 0 | 27.8 | 27.4 | 781 | 10.3 | 472 | 4 |
| -7 | MARV06 | 12.26 | 3.22 | 17.1 | 1.74 | 54 | 41.5 | 1.48 | 46 | 33.8 | 0 | 0 | 13.9 | 0 | 0 | 41.2 | 0 | 0 | 26.5 | 31.2 | 530 | 10.9 | 83.4 | 0.68 |
| 28 | MARV06 | 11.94 | 4.93 | 16.3 | 2.68 | 54.4 | 42.5 | 2.18 | 44.2 | 35.6 | 0.03 | 0.6 | 13.7 | 0 | 0 | 38.4 | 0.04 | 0.8 | 26 | 29.4 | 464 | 13.5 | 401.2 | 3.36 |
| -7 | MARV11 | 13.43 | 8.97 | 18.8 | 4.66 | 52 | 48.7 | 4.31 | 48 | 36.3 | 0 | 0 | 14 | 0 | 0 | 38.6 | 0 | 0 | 27.2 | 31 | 728 | 11.3 | 192 | 1.43 |
| 28 | MARV11 | 13.23 | 16.15 | 18 | 1.58 | 9.8 | 47.4 | 14.53 | 90 | 35.8 | 0.04 | 0.2 | 13.6 | 0 | 0 | 38 | 0 | 0 | 27.6 | 31.5 | 744 | 12 | 654.9 | 4.95 |
| -7 | MARV12 | 13.94 | 18.77 | 18.3 | 6.25 | 33.3 | 45.4 | 12.36 | 65.8 | 32.6 | 0.16 | 0.9 | 13.1 | 0 | 0 | 40.3 | 0 | 0 | 26.9 | 36 | 502 | 11.2 | 111.5 | 0.8 |
| 28 | MARV12 | 13 | 6.04 | 17.1 | 1.04 | 17.2 | 43.7 | 4.93 | 81.6 | 33.6 | 0.02 | 0.3 | 13.2 | 0.01 | 0.2 | 39.1 | 0.04 | 0.7 | 29.2 | 34.5 | 820 | 11.2 | 434.2 | 3.34 |

red blood cells (RBC), white blood cells (WBC), hemoglobin (HGB), neutrophil (NEUT), hematocrit (HCT), lymphocytes (LYMPH), mean corpuscular volume (MCV), monocytes (MONO), mean corpuscular hemoglobin (MCH), eosinophils (EOS), mean corpuscular hemoglobin concentration (MCHC), basophils (BASO), red cell distribution width (RDW-SD), red cell distribution width-coefficient of variation (RDW CV), platelets (PLT), mean platelet volume (MPV), reticulocytes (RET), number (#), percentage (%)

**Supplementary Table 3.** Clinical chemistry of healthy control and EBOV or MARV-infected JFBs. Blood urea nitrogen (BUN), alanine aminotransferase (ALT), alkaline phosphatase (ALP), aspartate aminotransferase (AST), sodium (Na+), potassium (K+), chlorine (Cl-), total carbon dioxide (tCO2), heme (Hem), lipase (Lip),

| Study Day | Animal ID | BUN (mg/dL) | Creatinine (mg/dL) | ALT (U/L) | ALP (U/L) | AST (U/L) | T Bilirubin (mg/dL) | Glucose (mg/dL) | Calcium (mg/dL) | T Protein (g/dL) | Albumin (g/dL) | Globulin (g/dL) | Na+ (mmol/L) | K+ (mmol/L) | Cl- (mmol/L) | tCO2 (mmol/L) | Hem | Lip | Notes |
| --- | --- | --- | --- | --- | --- | --- | --- | --- | --- | --- | --- | --- | --- | --- | --- | --- | --- | --- | --- |
| Healthy Control | BT01 | 9 | 0.4 | 83 | 106 | 188 | 0.3 | 53 | 9.5 | 6.4 | 3.0 | 3.4 | 145 | 3.8 | 108 | 23 | 0 | 0 |  |
| Healthy Control | BT02 | 9 | 0.2 | 64 | 139 | 106 | 0.2 | 130 | 10.5 | 6.0 | 3.1 | 2.9 | 146 | 4.6 | 106 | 22 | 0 | 0 |  |
| Healthy Control | BT03 | 9 | 0.4 | 66 | 101 | 235 | 0.2 | 175 | 9.9 | 6.4 | 3.4 | 3.0 | 138 | 3.9 | 103 | 24 | 0 | 0 |  |
| Healthy Control | BT04 | 10 | 0.3 | 69 | 98 | 128 | 0.3 | 157 | 9.5 | 6.5 | 3.4 | 3.1 | 144 | 4.8 | 106 | 26 | 0 | 0 |  |
| Healthy Control | BT05 | 4 | 0.4 | 76 | 146 | 125 | 0.2 | 76 | 10.2 | 6.2 | 3.2 | 3.0 | 145 | 3.9 | 100 | 24 | 0 | 0 |  |
| Healthy Control | BT06 | 6 | 0.2 | 80 | 76 | 193 | 0.2 | 61 | 9.4 | 5.8 | 2.9 | 2.9 | 143 | 4.6 | 108 | 23 | 0 | 0 |  |
| Healthy Control | BT07 | 10 | 0.6 | 135 | 300 | 283 | 0.2 | 89 | 9.9 | 5.5 | 2.7 | 2.8 | 145 | 5.9 | 113 | 24 | 0 | 0 |  |
| Healthy Control | BT08 | 7 | 0.3 | 65 | 72 | 172 | 0.2 | 158 | 9.9 | 6.1 | 2.6 | 3.5 | 138 | 4.3 | 106 | 22 | 0 | 0 |  |
| Healthy Control | BT09 | 5 | 0.2 | 51 | 48 | 179 | 0.2 | 134 | 9.1 | 5.6 | 2.4 | 3.2 | 142 | 5.1 | 110 | 18 | 0 | 0 |  |
| Healthy Control | BT10 | 5 | <0.2 | 96 | 128 | 244 | 0.2 | 151 | 10.0 | 5.6 | 2.9 | 2.7 | 139 | 5.5 | 105 | 20 | 0 | 0 |  |
| 3 | EBO01 | 7 | 0.4 | 85 | 60 | 306 | 0.2 | 126 | 8.8 | 5.8 | 3.1 | 2.8 | 150.0 | 6.3 | 105 | 23 | 2 | 0 |  |
| 3 | EBO02 | 9 | 0.2 | 103 | 106 | 238 | 0.2 | 138 | 9.6 | 6.0 | 3.1 | 2.8 | 151 | 8.5 | 107 | 16 | 0 | 0 |  |
| 3 | EBO07 | 11 | <0.2 | 55 | 83 | 173 | 0.3 | 111 | 8.1 | 5.8 | 3.1 | 2.7 | 146 | 6.2 | 103 | 26 | 0 | 0 |  |
| 7 | EBO08 | 8 | 0.2 | 54 | 72 | 115 | 0.2 | 160 | 9.3 | 6.2 | 3.1 | 3.1 | 151.0 | 7.4 | 107 | 22 | 0 | 0 |  |
| 7 | EBO03 |  |  |  |  |  |  |  |  |  |  |  |  |  |  |  |  |  | Insufficient sample |
| 7 | EBO04 |  |  |  |  |  |  |  |  |  |  |  |  |  |  |  |  |  | Insufficient sample |
| 7 | EBO09 | 8 | 0.3 | 65 | 71 | 122 | 0.2 | 290 | 8.6 | 6.0 | 3.0 | 3.1 | 142 | >8.5 | 99 | 14 | 0 | 0 |  |
| 28 | EBO10 | 8 | <0.2 | 59 | 84 | 111 | 0.1 | 200 | 10.5 | 6.9 | 3.4 | 3.5 | 150.0 | 8.4 | 107 | 22 | 0 | 0 |  |
| 28 | EBO05 |  |  |  |  |  |  |  |  |  |  |  |  |  |  |  |  |  | Insufficient sample |
| 28 | EBO06 | 8 | <0.2 | 51 | 78 | 98 | 0.2 | 191 | 9.3 | 6.1 | 3.0 | 3.1 | 144 | 6.2 | 107 | 26 | 0 | 0 |  |
| 28 | EBO11 | 7 | 0.2 | 37 | 65 | 81 | <0.1 | 226 | 10.7 | 6.3 | 3.3 | 3.0 | 145 | >8.5 | 108 | 25 | 0 | 0 |  |
| 3 | EBO12 |  |  |  |  |  |  |  |  |  |  |  |  |  |  |  |  |  | Insufficient sample |
| 3 | MARV01 |  |  |  |  |  |  |  |  |  |  |  |  |  |  |  |  |  | Insufficient sample |
| 3 | MARV02 | 7 | <0.2 | 71 | 106 | 397 | 0.2 | 140 | 9.3 | 6.0 | 3.0 | 3.0 | 146 | 6.2 | 106 | 18 | 1 | 0 |  |
| 3 | MARV07 | 12 | <0.2 | 87 | 65 | 191 | 0.2 | 150 | 9.4 | 5.8 | 3.1 | 2.7 | 143 | 6.0 | 101 | 23 | 1 | 0 |  |
| 7 | MARV08 | 8 | 0.2 | 61 | 67 | 149 | 0.2 | 202 | 8.6 | 6.1 | 3.2 | 2.9 | 142 | 5.8 | 103 | 20 | 0 | 0 |  |
| 7 | MARV03 | 7 | 0.4 | 46 | 56 | 135 | 0.3 | 189 | 9.4 | 6.7 | 3.6 | 3.0 | 145 | 7.6 | 100 | 21 | 0 | 0 |  |
| 7 | MARV04 | 7 | 0.3 | 44 | 97 | 102 | 0.2 | 305 | 10.3 | 5.9 | 3.0 | 2.9 | 146 | >8.5 | 97 | 23 | 0 | 0 |  |
| 7 | MARV09 | 10 | <0.2 | 43 | 67 | 90 | 0.2 | 192 | 9.7 | 6.4 | 3.6 | 2.8 | 149 | >8.5 | 104 | 19 | 2 | 0 |  |
| 28 | MARV10 |  |  |  |  |  |  |  |  |  |  |  |  |  |  |  |  |  | Insufficient sample |
| 28 | MARV05 | 6 | 0.3 | 55 | 97 | 132 | 0.1 | 243 | 10.4 | 6.5 | 3.3 | 3.2 | 147 | >8.5 | 105 | 18 | 2 | 0 |  |
| 28 | MARV06 | 8 | 0.2 | 48 | 115 | 201 | 0.1 | 347 | 10.6 | 5.9 | 2.9 | 3.0 | 145 | >8.5 | 103 | 24 | 0 | 0 |  |
| 28 | MARV11 | 10 | <0.2 | 71 | 67 | 113 | 0.1 | 163 | 9.1 | 6.1 | 3.2 | 2.9 | 146 | 6.7 | 107 | 24 | 0 | 0 |  |

**Supplementary Table 4.** Differentially Upreglated genes in EBOV or MARV-infected JFBs compared to healthy controls.

| Challenge Virus | Direction | adj.Pval | nGenes | Pathways |
| --- | --- | --- | --- | --- |
| EBOV | Up regulated | 9.30E-63 | 176 | Immune response |
|  | Up regulated | 1.19E-56 | 199 | Immune system process |
|  | Up regulated | 7.37E-50 | 136 | Defense response |
|  | Up regulated | 2.30E-42 | 121 | Response to biotic stimulus |
|  | Up regulated | 8.84E-42 | 118 | Response to other organism |
|  | Up regulated | 2.03E-40 | 124 | Biological process involved in interspecies interaction between organisms |
|  | Up regulated | 8.79E-40 | 100 | Defense response to other organism |
|  | Up regulated | 1.26E-39 | 91 | Innate immune response |
|  | Up regulated | 1.28E-38 | 125 | Regulation of immune system process |
| MARV | Up regulated | 4.47E-27 | 58 | Immune response |
|  | Up regulated | 4.23E-26 | 50 | Defense response |
|  | Up regulated | 2.85E-25 | 64 | Immune system process |
|  | Up regulated | 3.42E-24 | 48 | Biological process involved in interspecies interaction between organisms |
|  | Up regulated | 4.25E-23 | 37 | Innate immune response |
|  | Up regulated | 5.13E-23 | 45 | Response to biotic stimulus |
|  | Up regulated | 9.83E-23 | 44 | Response to other organism |
|  | Up regulated | 1.78E-22 | 39 | Defense response to other organism |
|  | Up regulated | 1.62E-18 | 43 | Regulation of immune system process |

**Supplementary Table 5.** Differentially Upreglated genes in EBOV-infected compared to MARV-infected JFBs.

| Direction | adj.Pval | nGenes | Pathways | Genes |
| --- | --- | --- | --- | --- |
| Up regulated | 0.000110<br>27 | 4 | Negative regulation of viral process | RSAD2 PML ISG15 LY6E |
| Up regulated | 0.000110<br>27 | 4 | Type I interferon signaling pathway | RSAD2 ISG15 PTPN1 PSMB8 |
| Up regulated | 0.000110<br>27 | 4 | Cellular response to type I interferon | RSAD2 ISG15 PTPN1 PSMB8 |
| Up regulated | 0.000460<br>32 | 2 | Cytosol to endoplasmic reticulum transport | TAP2 TAP1 |
| Up regulated | 0.000460<br>32 | 4 | Regulation of viral life cycle | RSAD2 PML ISG15 LY6E |
| Up regulated | 0.000846<br>493 | 5 | Regulation of response to biotic stimulus | IL4I1 PML ISG15 PTPN1 PSMB8 |
| Up regulated | 0.000846<br>493 | 9 | Immune response | IL4I1 RSAD2 IFI44L PML ISG15 PTPN1 TAP2 PSMB8 TAP1 |
| Up regulated | 0.000846<br>493 | 4 | Regulation of viral process | RSAD2 PML ISG15 LY6E |
| Up regulated | 0.000846<br>493 | 2 | Vesicle fusion with endoplasmic reticulum-Golgi intermediate compartment (ERGIC) membrane | TAP2 TAP1 |
| Up regulated | 0.000939<br>666 | 7 | Viral process | RSAD2 PML ISG15 TAP2 PSMB8 TAP1 LY6E |
| Up regulated | 0.001179<br>496 | 3 | Antigen processing and presentation of exogenous peptide antigen via MHC class I, TAP-dependent | TAP2 PSMB8 TAP1 |
| Up regulated | 0.001179<br>496 | 2 | Antigen processing and presentation of endogenous peptide antigen | TAP2 TAP1 |
| Up regulated | 0.001201<br>765 | 4 | Defense response to virus | RSAD2 IFI44L PML ISG15 |
| Up regulated | 0.001458<br>967 | 7 | Response to biotic stimulus | IL4I1 RSAD2 IFI44L PML ISG15 PTPN1 PSMB8 |
| Up regulated | 0.001905<br>862 | 6 | Defense response to other organism | RSAD2 IFI44L PML ISG15 PTPN1 PSMB8 |

**Supplementary Figure 1.**
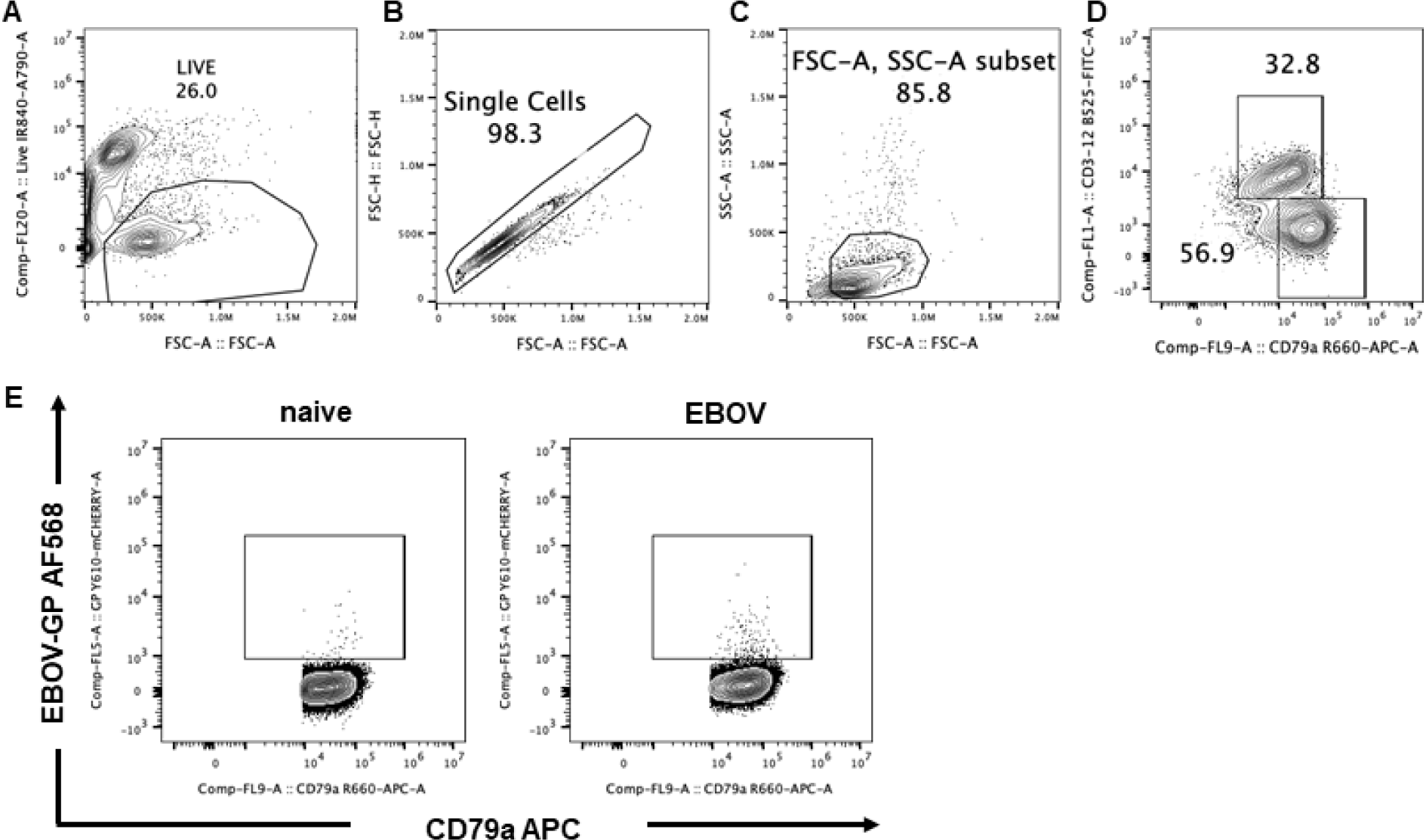
Representative gating strategy for flow cytometric analysis of splenocytes. (A) Splenocytes were first gated on FSC-A vs the Live-Dead Stain to exclude dead cells. (B) Live cells were then gated to single cells using FSC-A vs FSC-H. (C) A lymphocyte gate utilizing FSC-A vs SSC-A was applied to live, single cells. (D) The live lymphocyte population was further analyzed by gating on B cells and T cells using intracellular staining of anti-CD79a APC vs anti-CD3 FITC, respectively. (E) Representative FACS contour plots showing the background staining (naïve) and staining (EBOV) of Alexa-fluor 568 conjugated recombinant EBOV GP receptor binding domain on splenic B cells

**Supplementary Figure 2.**
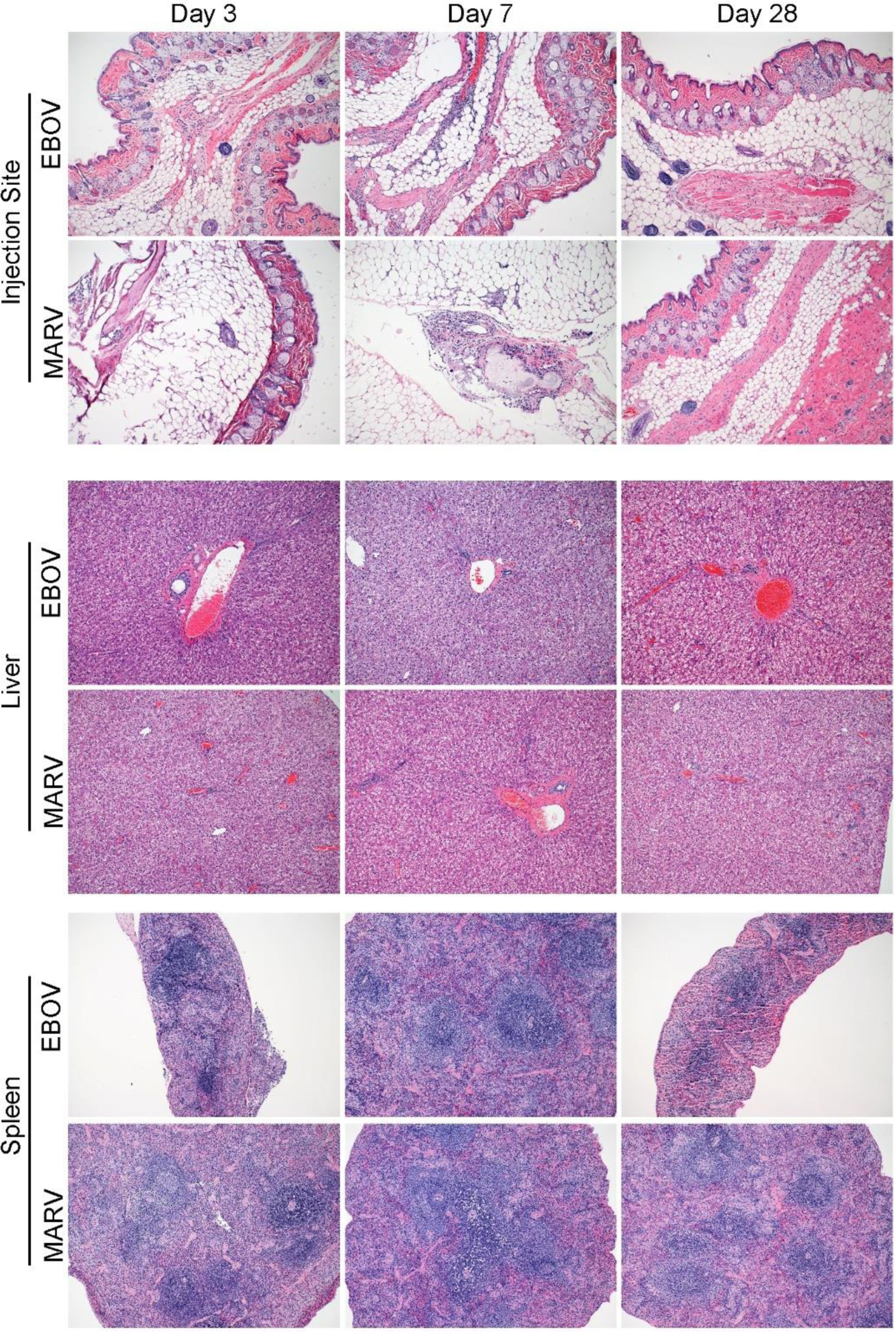
Infection with EBOV or MARV does not induce histopathologic changes in JFBs. Tissue sections were stained with hematoxylin and eosin. Representative sections for skin at the injection site, liver, and spleen sections for Ebola virus (EBOV) or Marburg virus (MARV) challenged bats at 3, 7, and 28 days post-inoculation. The following magnification was used for all slides: 100X.

**Supplementary Figure 3.**
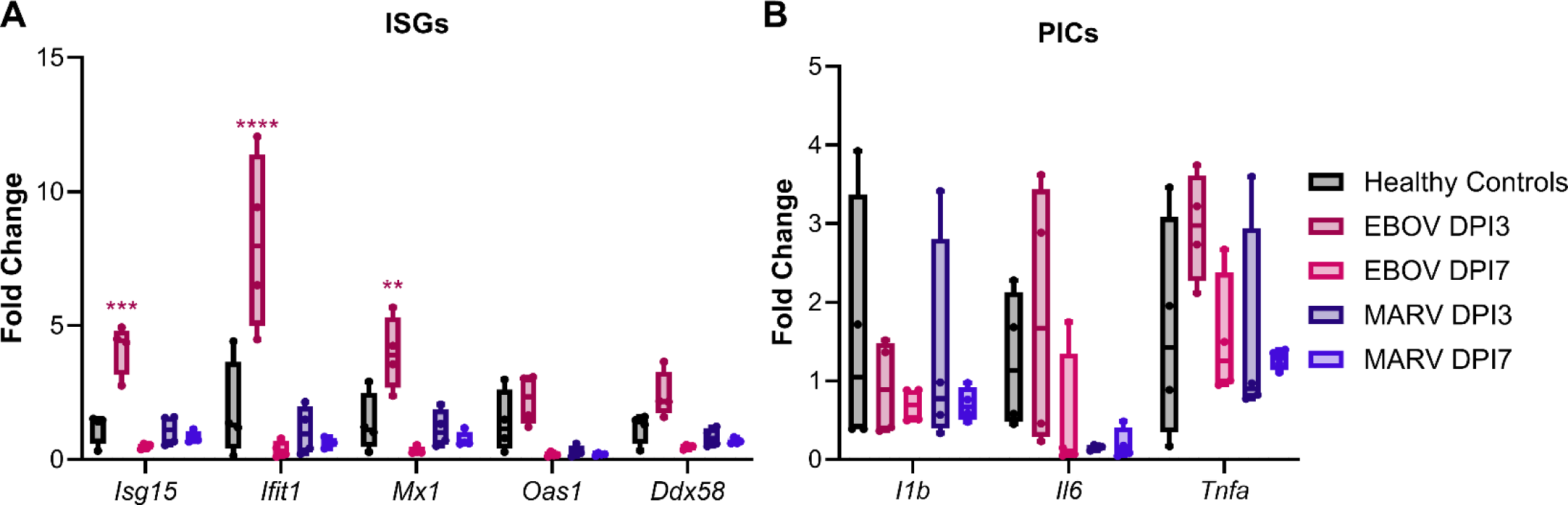
EBOV infection induces mild ISG expression in the lung. (A) RT-qPCR of interferon-stimulate genes (ISGs) mRNA in lungs collected from healthy control bats (n=4) and at day (D) 3 or 7 necropsy of EBOV- or MARV-infected bats (n=4). (B) RT-qPCR of pro-inflammatory cytokines (PICs) mRNA in lungs collected from healthy control bats (n=4) and at day (D) 3 or 7 necropsy of EBOV- or MARV-infected bats (n=4). (A-B) ΔC_T_ values were normalized to the average ΔC_T_ value of the healthy control bats to calculate ΔΔC_T_. Fold change of each gene at necropsy day was compared to the healthy controls using a two-way ANOVA with Dunnett’s multiple comparison.

**Supplementary Figure 4.**
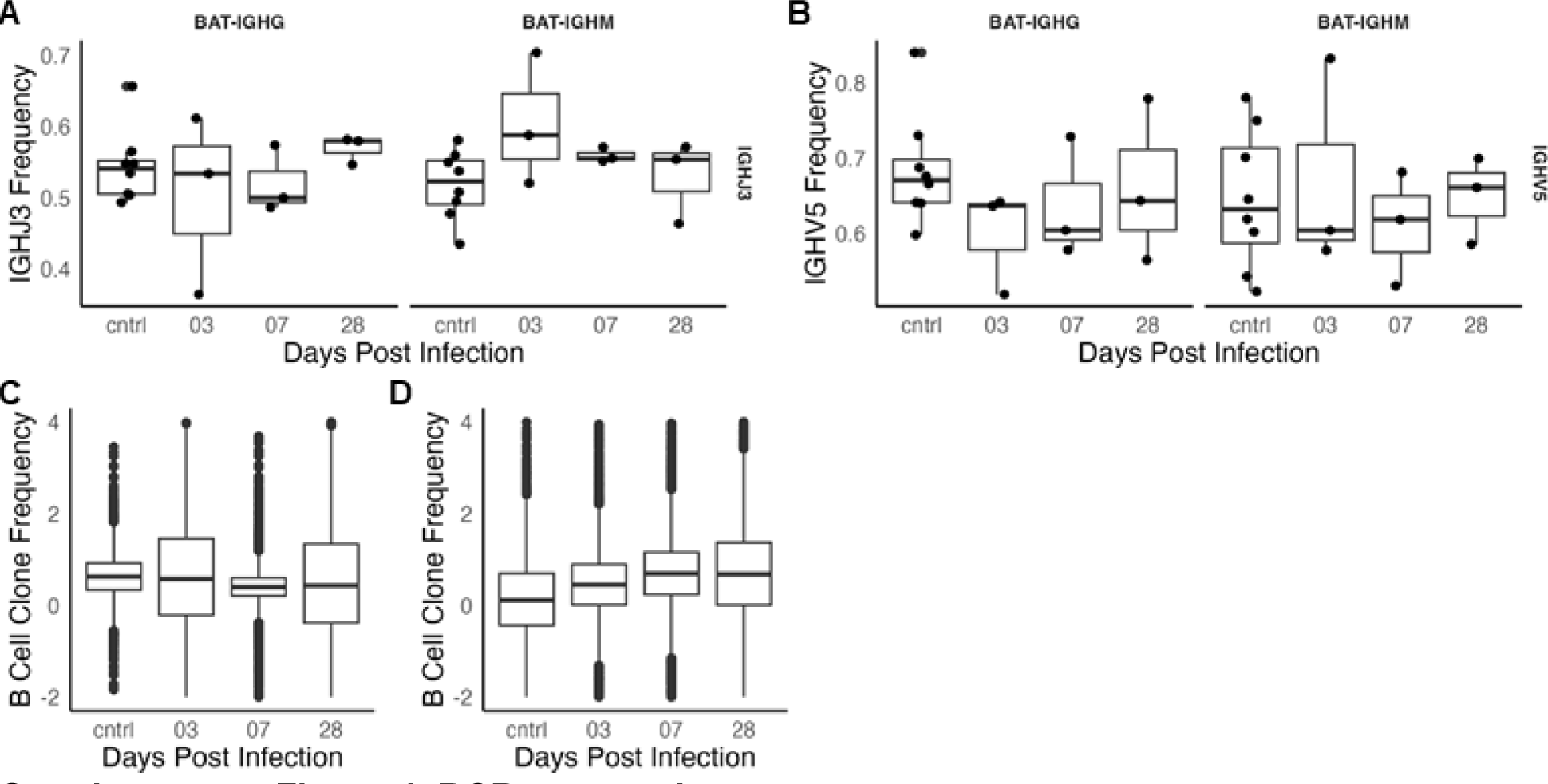
BCR sequencing. (A) Clonal frequencies of IgG sequences in the spleens of healthy control bats (n=8) or Ebola virus challenged bats at day 3, 7, and 28 necropsies (n=4). (B) Clonal frequencies of IgM BCR sequences. (A-B) Data represents a sample from the posterior effect estimates obtained from a Bayesian hierarchical model. Points represent the frequency observed in individual bats. Boxplots show the 25th, 50th (median), and 75th percentiles, with lines indicating the smallest and largest values within 1.5 times the interquartile range. (C) Frequency of BCR sequences containing the putative IGHV5 gene segment. Dots represent data from individual bats. (D) Frequency of BCR sequences containing the putative IGHJ3 gene segment. Dots represent data from individual bats.

**Supplementary Figure 5.**
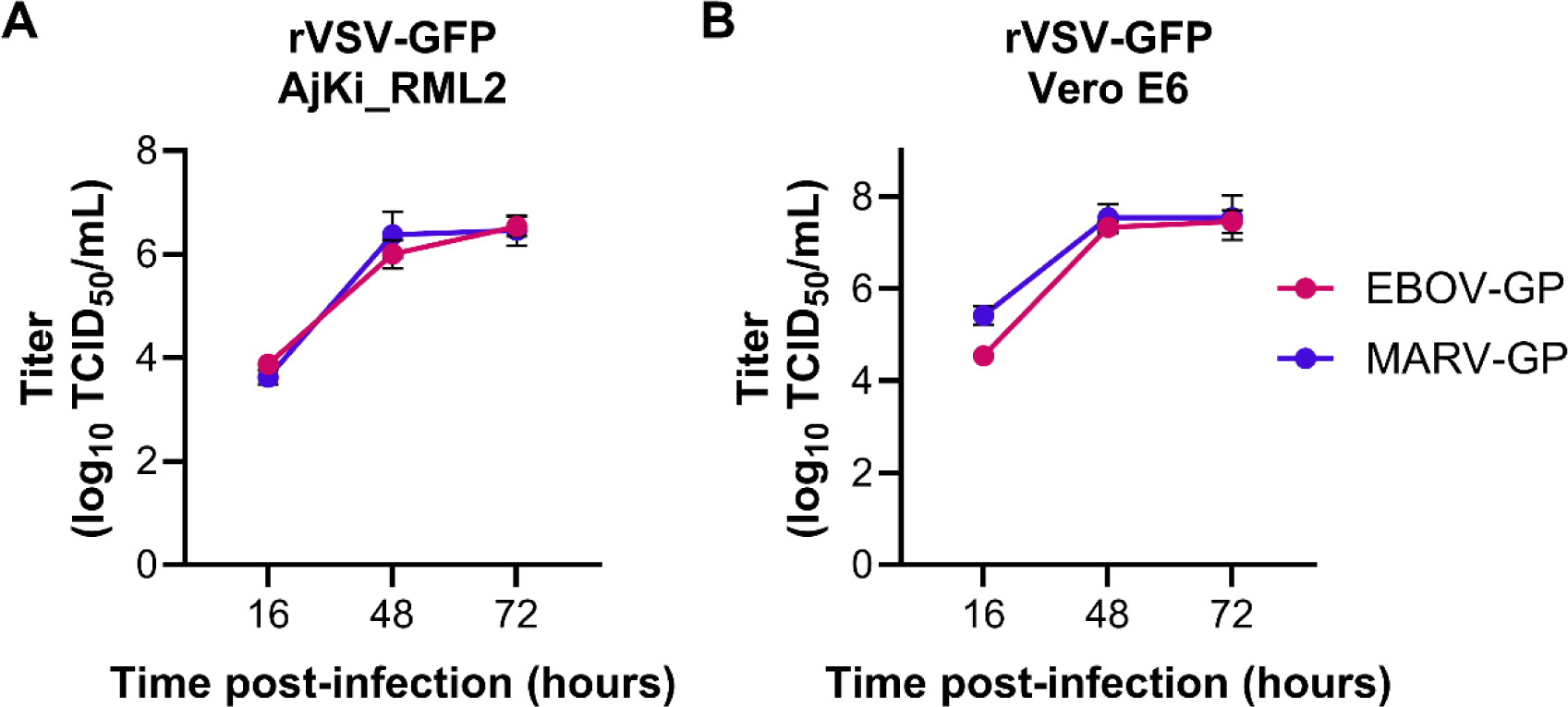
Replication kinetics of recombinant VSV expressing EBOV or MARV GP. (A) Infectious titers of rVSV-EBOV-GFP or rVSV-MARV-GFP from AjKi-RML2 cells infected at multiplicity of infection (MOI) 0.005 at 16, 48, and 72 hours post-infection. (B) Infectious titers of rVSV-EBOV-GFP or rVSV-MARV-GFP from Vero E6 cells infected at multiplicity of infection (MOI) 0.005 at 16, 48, and 72 hours post-infection. (A-B) Two independent experiments were performed with 3 technical replicates for each experiment. The titer at each time point was compared using a two-way ANOVA with Dunnett’s multiple comparison.

**Supplementary Figure 6.**
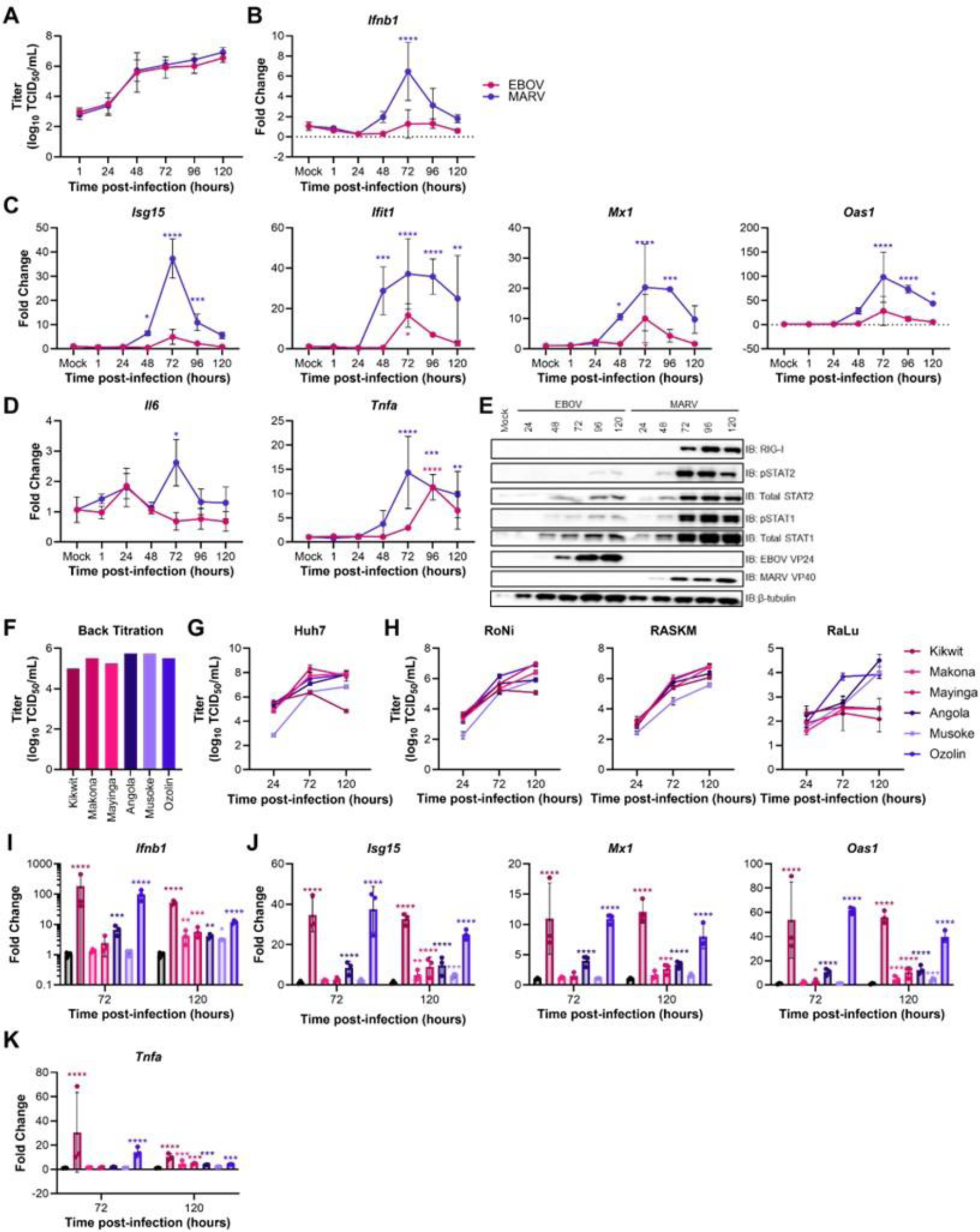
EBOV and MARV antagonize the IFN-I response with similar efficiencies in ERB cells. (A) Infectious titers of EBOV-Mayinga or MARV-Ozolin on Egyptian rousette bat kidney cells (RoNi) infected with multiplicity of infection (MOI) 0.1. Data presented as log_10_ transformed values. Data is from two independent experiments with three technical replicates per experiment. Two-way ANOVA with Sidak’s multiple test comparison was performed to evaluate the difference between EBOV and MARV titer at each time point. (B) RT-qPCR of *Ifnb1* mRNA in RoNi cells infected with EBOV or MARV at MOI 0.1. (C) RT-qPCR of interferon stimulated gene mRNAs in RoNi cells infected with EBOV or MARV at MOI 0.1. (D) RT-qPCR of pro-inflammatory cytokine mRNAs in RoNi cells infected with EBOV or MARV at MOI 0.1. (E) Immunoblot of RoNi cells infected with EBOV or MARV at MOI 0.1. The presented panel is representative of three independent immunoblots. (F) Infectious titers of inoculum of the three EBOV and MARV strains used to infect Huh7 (S5G), AjKi_RML2 (6F), and Egyptian rousette cells (S5H) to assess replication kinetics. (G) Infectious titers of three EBOV strains and three MARV strains on human hepatoma (Huh7), cells infected with MOI 0.1. Data presented as log_10_ transformed values. Data is from one experiment conducted in triplicate. (H) Infectious titers of three EBOV strains and three MARV strains on Egyptian rousette bat immortalize kidney (RoNi), primary kidney (RASKM), and primary lung (RaLu) cells infected with MOI 0.1. Data presented as log_10_ transformed values. Data is from one experiment conducted in triplicate. (I) RT-qPCR of *Ifnb1* mRNA in RoNi cells infected with three different EBOV or MARV strains at MOI 0.1. (J) RT-qPCR of ISG mRNAs in RoNi cells infected with three different EBOV or MARV strains at MOI 0.1. (K) RT-qPCR of pro-inflammatory cytokine *Tnfa* mRNAs in RoNi cells infected with three different EBOV or MARV strains at MOI 0.1. (B-D;I-K) ΔC_T_ values were normalized to the average ΔC_T_ value of mock infected cells to calculate ΔΔC_T_. Fold change of each gene at each time point for EBOV and MARV-infected cells was compared mock using a two-way ANOVA with Dunnett’s multiple test comparison.

**Supplementary Figure 7.**
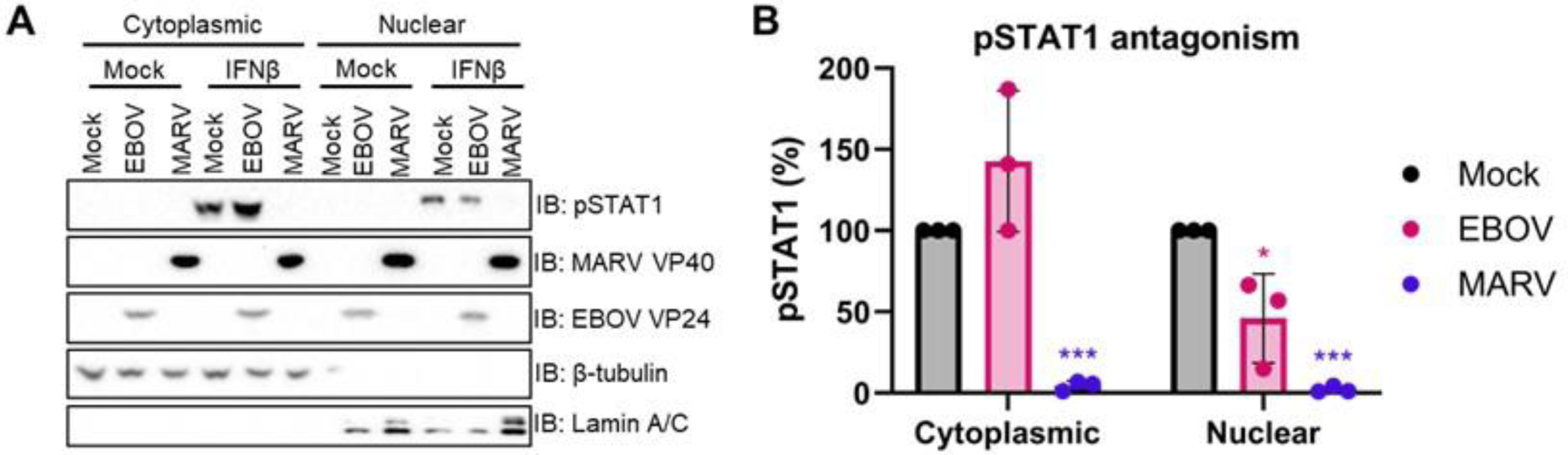
EBOV and MARV efficiently antagonize human IFN-I signaling. (A) Immunoblot of cytoplasmic and nuclear fractions of human hepatoma (Huh7) cells infected with MOI 1.5 of EBOV-Mayinga or MARV-Ozolin for 24 hours prior to the addition of exogenous recombinant Chiroptera IFN-β (100ng/mL) for 30 minutes. The presented panel is representative of three independent immunoblots. (B) Quantification of phosphorylated STAT1 in the cytoplasmic and nuclear fractions. Normalized expression was determined using the following formula: (AUC pSTAT1)/(AUC β-tubulin (cytoplasmic) or Lamin A/C (nuclear)). To calculate relative expression, the normalized sample value is divided by the normalized expression in uninfected cells treated with IFN-β. Three independent experiments were performed. Percentage of pSTAT1 in each of the fractions for EBOV- and MARV-infected cells was compared mock using a two-way ANOVA with Dunnett’s multiple test comparison.

**Supplementary Figure 8.**
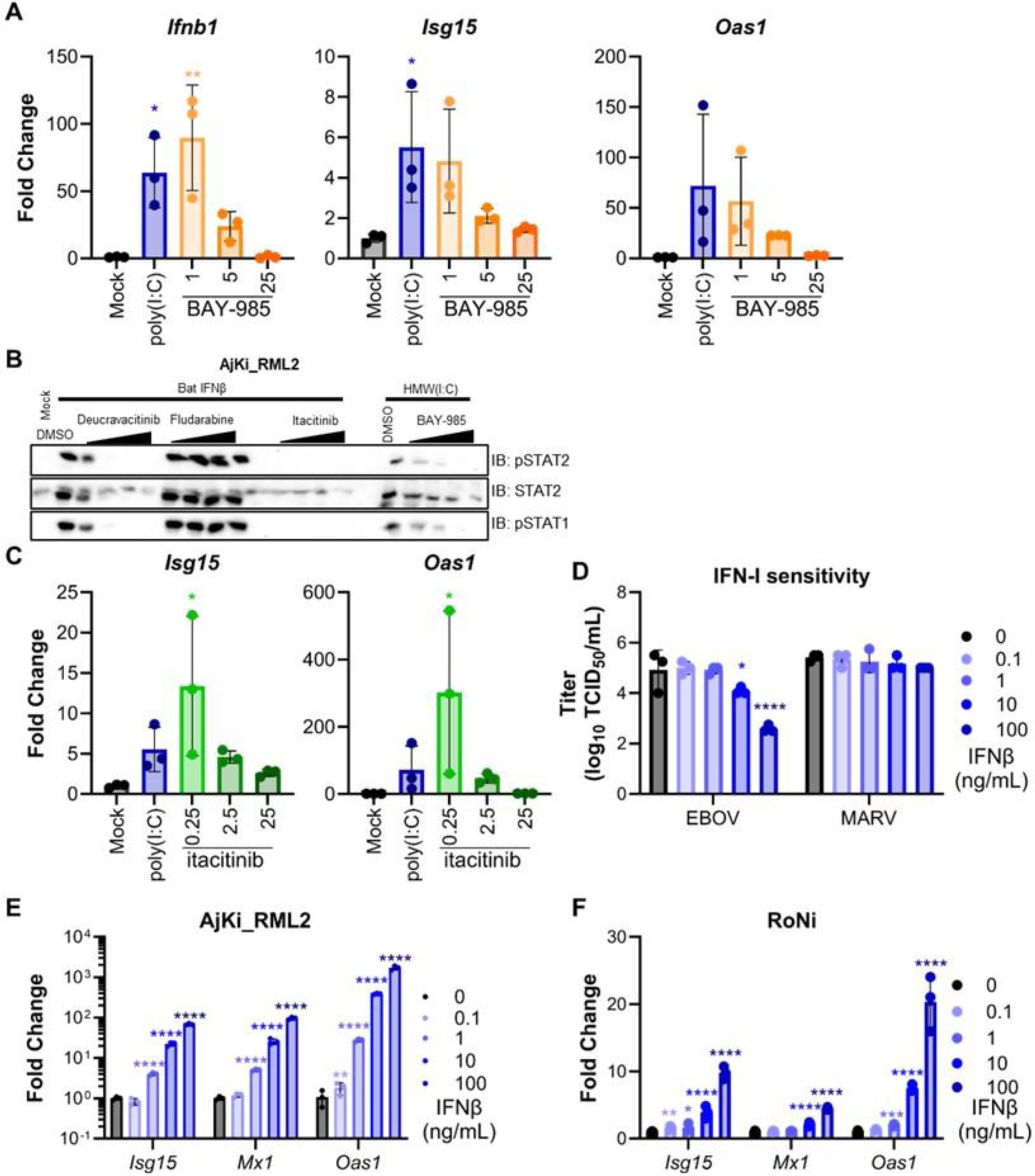
Validation of Chiroptera IFN-β and IFN-I inhibitors in JFB cells. (A) RT-qPCR of *Ifnb1* or interferon-stimulate gene (ISG), *Isg15* and *Oas1*, mRNAs in Jamaican fruit bat kidney cells, AjKi_RML2, were treated with type I interferon induction inhibitor BAY-985 (1-25 nM) for 24 hours prior to transfection of high molecular weight poly(I:C) for 18 hours. (B) Immunoblot of AjKi_RML2 cells treated with IFN-I signaling (deucravacitinib, fludarabine, or itacitinib) or induction (BAY-985) inhibitors at 5-fold dilution series (0.064-40nM) or treated with vehicle control, DMSO, for 24 hours prior to addition of recombinant Chiroptera IFN-β (100ng/mL) for 30 minutes (IFN-I signaling inhibitors) or transfection of high molecular weight poly(I:C) (500 ng/mL) for 24 hours (BAY-985). (C) RT-qPCR of *Isg15* and *Oas1*, mRNAs in AjKi_RML2 cells treated with type I signaling inhibitor itacitinib (0.25-25 nM) for 24 hours prior to transfection of high molecular weight poly(I:C) for 18 hours. (A-C) ΔC_T_ values were normalized to the average ΔC_T_ value of mock infected cells to calculate ΔΔC_T_. Fold change of each gene at dose was compared to unstimulated cells treated with DMSO using a one-way ANOVA with Dunnett’s multiple test comparison. (D) Infectious titers of EBOV-Mayinga or MARV-Ozolin at 48 hours post-infection. RoNi cells were treated with recombinant Chiroptera IFN-β in 10-fold serial dilution (0-100 ng/mL) for 24 hours prior to infection with MOI 0.1. Data presented as log_10_ transformed values. The experiment was performed in triplicate. To test a change in viral titer at each dose compared to mock treated cells, we performed a two-way ANOVA with Dunnett’s multiple test comparison. (E) RT-qPCR of interferon-stimulate gene, ISG15, Mx1, and OAS1, mRNAs in AjKi_RML2 cells treated with 10-fold serial concentrations of recombinant Chiroptera IFN-β for 24 hours. (F) RT-qPCR of interferon-stimulate gene, ISG15, Mx1, and OAS1, mRNAs in RoNi cells treated with 10-fold serial concentrations of recombinant Chiroptera IFN-β for 24 hours. (E-F) ΔΔC_T_ values (n=3) were normalized to the average mock ΔΔC_T_ value, and the data was log_10_ transformed. Fold change of each gene at IFN-β dose was compared to mock using a two-way ANOVA with Dunnett’s multiple test comparison.

